# Genomic rDNA instabilities in Arabidopsis epigenetic mutants alter location-based rRNA gene expression patterns

**DOI:** 10.64898/2026.03.11.710966

**Authors:** Murali Krishna Ramgopal, Abirami T. Subramanian, Ramya Tammineni, Aveepsha Bera, Balamurugan Aravind, Snigdha Ghosh, Gargi P. Saradadevi, Sahana Ravi, Gireesha Mohannath

## Abstract

The model plant species *Arabidopsis thaliana* has two nucleolus organizer regions (NORs), each consisting of tandem repeats of 45S rRNA genes on chromosomes 2 (*NOR2*) and 4 (*NOR4*). In the ecotype Col-0, the rRNA gene subtypes mapping to *NOR2* are mostly silenced during development, whereas those mapping to *NOR4* are mostly active. Using molecular and genetic mapping approaches, we demonstrate in multiple epigenetic mutants the occurrence of NOR conversions involving loss of rDNA and the associated telomeres in one NOR and replacement of the lost sequences with the corresponding sequences from the other NOR. We studied mutants of evolutionarily conserved chromatin remodelers *DECREASE IN DNA METHYLATION 1* (*DDM1*) and *CHROMATIN ASSEMBLY FACTOR 1* (*CAF-1*), and plant-specific *CHROMOMETHYLASE 2* (*CMT2*) DNA methyltransferase. We observed NOR conversions in *ddm1* and *caf-1* mutants, where such NOR conversions altered the rRNA variant expression patterns, which often confound the data pertaining to release of rRNA gene silencing. We delineated the effects of these mutations on the rDNA instability-mediated alterations in rRNA variant expression from their effects on the actual release of rRNA gene silencing. We also show that these mutations release rRNA gene silencing independently of their effect on rDNA genomic instability involving NOR conversions.

## Introduction

In eukaryotes, hundreds to thousands of ribosomal RNA (rRNA) genes are tandemly arrayed at chromosomal locations called nucleolus organizer regions (NORs). These rRNA genes are transcribed in the nucleolus by RNA polymerase I, yielding primary transcripts, which vary in size from 35S to 48S depending on the species. These transcripts are processed into 18S, 5.8S, and 25S–28S rRNAs that constitute the catalytic cores of ribosomes, the protein-synthesizing cell organelles (1–4). Model plant species *Arabidopsis thaliana* has two NORs; one each on the shorter arms of chromosomes 2 (*NOR2*) and 4 (*NOR4*) (5, 6). In the ecotype Columbia-0 (Col-0), *NOR2* and *NOR4* comprise ∼5.5– and ∼3.9 Mb of rRNA gene repeats, respectively, with each rRNA gene spanning ∼10 kb (7).

All rRNA genes are nearly identical, but length and sequence variation exist among them. Based on the existing polymorphism in the 3’ ETS region, four rRNA gene subtypes were described in Col-0, numbered Variant1 through 4 (VAR1 to VAR4) (8–10). A recent study in Col-0 ecotype involving *de novo* assembly of NORs identified a total of 74 rRNA gene subtypes, based on length and sequence variation (7).

The rRNA gene expression is developmentally regulated as a gene dosage control mechanism, which in Col-0 has been demonstrated to operate at the level of NORs, wherein *NOR2* genes undergo selective silencing and the *NOR4* genes remain transcriptionally active (8, 9). Relatively, *NOR2* genes are hypermethylated and exist in more condensed chromatin when compared to *NOR4* genes (7, 11–13). Thus, the distinct rRNA subtypes and their association with separate NORs have rendered Col-0 an ideal Arabidopsis ecotype for studying mechanisms of selective rRNA gene silencing.

In an earlier study involving Arabidopsis *atxr5 atxr6* mutants, defective for histone 3 lysine 27 monomethylation (H3K27me1), abundant expression of VAR1 had led to the conclusion that the loss of ATXR5 and ATXR6 significantly impacts *NOR2* gene silencing (14). However, a later study employing genetic linkage-based mapping of rRNA variants and Fluorescence-Activated Nucleolus Sorting (FANoS), demonstrated that in the *atxr5 atxr6* mutants, a multimegabase-scale loss of 45S rDNA and the associated telomere of *NOR4* and its repair from the corresponding *NOR2* sequences has occurred (9). Indeed, this translocation of VAR1 genes from *NOR2* to *NOR4* led to their expression on *NOR4*, while still remaining silent on *NOR2* (9). Further, the study revealed that the effect of *atxr5/6* mutations on the release of *NOR2* gene silencing was rather modest (9). Collectively, this study revealed that the mere abundant expression of VAR1 genes in Col-0 mutants does not necessarily indicate the release of *NOR2* gene silencing. Therefore, we studied multiple Arabidopsis epigenetic mutants that are defective in chromatin condensation and DNA methylation, to determine their effect on rDNA genomic instability and whether such genomic instabilities would affect rRNA gene expression pattern.

In Arabidopsis, *METHYLTRANSFERASE 1 (MET1)* and *CHROMOMETHYLASE 3 (CMT3)* mediate virtually all of CG and CHG methylation (where H denotes A, T, or C), respectively (15–17). CHH methylation is directed by small RNAs and mediated by *DOMAINS REARRANGED METHYLTRANSFERASE 2* (*DRM2*) via the RNA-directed DNA methylation (RdDM) pathway, but also by *CHROMOMETHYLASE 2* (*CMT2*) independently of RdDM (17–20). A recent study has indicated that most of the CHH methylation at rDNA loci is carried out by CMT2, and its deficiency led to significant disruption of *NOR2* gene silencing (13). CMT2 and CMT3 belong to a group of plant-specific chromomethylases (21). *DECREASE IN DNA METHYLATION 1* (*DDM1*), a SWI2/SNF2 family chromatin remodeler, acts as a master regulator of DNA methylation (22–24). Loss of DDM1 results in genome-wide demethylation at 70% of methylated cytosines in all three sequence contexts (CG, CHG, and CHH) (17, 25, 26), and also disrupts *NOR2* gene silencing (13).

*CHROMATIN ASSEMBLY FACTOR 1* (*CAF-1*) is an evolutionarily conserved heterotrimeric histone chaperone that plays a vital role in depositing histone H3.1 and H4, whereas other histone chaperones deposit histone H2A and H2B during the DNA synthesis process (27–29). In *Arabidopsis thaliana*, the three subunits are encoded by the *FASCIATA1* (*FAS1*), *FAS2*, and *MULTICOPY SUPPRESSOR OF IRA1* (*MSI1*) genes (30–32). Loss of CAF-1 led to successive loss of rDNA over generations and disruption of *NOR2* gene silencing (33, 34). Note that, ATXR5 and ATXR6 monomethylate histone H3.1, which CAF-1 deposits to the chromatin (35). Therefore, we hypothesize that one of the reasons for the disruption of *NOR2* gene silencing in *caf-1* mutants might be due to the translocation of rRNA genes from *NOR2* to *NOR4*.

We also hypothesized that similar NOR conversion-mediated altered rRNA variant expression could have occurred in *ddm1* and *cmt2* mutants. To test these hypotheses, we studied multiple mutant alleles of *DDM1* (*ddm1-1* and *ddm1-2*), *CAF-1* (*fas1-4* and *fas2-4*), and *CMT2* (*cmt2-3*) to determine their effects on rDNA genomic stability and rRNA variant expression patterns. We chose *ddm1* and *cmt2* mutants because they significantly disrupt *NOR2* gene silencing (13). Previous studies have demonstrated a similar *NOR2* gene silencing in *caf-1* mutants (33, 34), but it remains to be investigated if rDNA instability found in these mutants involves translocation of rRNA genes from one NOR to the other, and how such a translocation would affect rRNA gene expression pattern. In this study, we demonstrate that translocation of rRNA genes between NORs has occurred in *ddm1* and *caf-1* mutants, but not in *cmt2* mutants. Translocation of rRNA genes from one NOR to the other in *ddm1* and *caf-1* mutants affects rRNA gene variant expression patterns. We also demonstrate that *NOR2* gene silencing is disrupted in all these mutants, independent of rDNA genomic instability.

## Materials and Methods

### Plant materials and growth conditions

Seed stocks of *Arabidopsis thaliana* ecotypes Col-0 (stock no. CS22681), Sha (stock no. CS22690), *cmt-3*, *ddm1-1*, *ddm1-2*, *fas1-4*, *and fas2-4* are gifts from the Pikaard Lab (Indiana University). All the mutants used in this study were generated in the Col-0 background. The genetic crosses were performed using the mutants or Col-0 as the pollen donors and the Sha ecotype as the female parent. The F_1_ plants from these crosses were allowed for self-fertilization to obtain F_2_ progenies, which were used for genetic mapping of rRNA variants and other NOR-associated markers. All plants were grown in soil media under controlled conditions with a 16-h light/8-h dark photoperiod cycle, 50–55% humidity, and a temperature of 23-25°C.

### DNA Extraction

Four-week-old leaf tissues were used to isolate DNA using a modified protocol from Edwards et al. (1991). Tissues were snap frozen, powdered, and suspended in a lysis solution (containing 200 mM Tris-HCl (pH 8.0), 250 mM NaCl, 2.5 mM EDTA pH 8.0), and 1% SDS), followed by centrifugation at 13000 g for 10 minutes at room temperature. The supernatant was then transferred to a new microcentrifuge tube containing one volume of phenol: chloroform: isoamyl alcohol (25:24:1). The mixture was then mixed by inverting, followed by centrifugation at 13000 g. The aqueous phase was collected in a new tube, and 3M sodium acetate (pH 5.2), equivalent to 1/10^th^ the volume of lysis buffer, and one volume of absolute ethanol were added and mixed properly. The mixture was then centrifuged at 13000 g at RT, and the supernatant was discarded. Next, the pellet was washed with 70% ethanol and kept for drying until all the ethanol evaporated. The dried pellet was then dissolved in Tris-EDTA (10mM Tris-Cl (pH 8.0), 0.25 mM EDTA (pH 8.0) buffer containing RNase A (20 µg/mL) and incubated for 30 minutes at 37 °C. The DNA samples were used for PCR assays, either with or without dilution, depending on the DNA concentration.

### Genotyping and PCR assays

For genotyping *ddm1-1* and *ddm1-2* mutants, which carry single-base substitutions, we used previously established CAPS or dCAPS assays that involve PCR amplification of the region carrying the mutation, followed by digestion with the respective restriction enzymes. The PCR amplicons are digested in the wild-type or mutants samples, depending on the presence or absence of the restriction site. For both *ddm1-1* and *ddm1-2* genotyping, PCR amplification was carried out using CAPS or dCAPS assays. Primers, PCR conditions, and restriction enzymes, used for CAPS-dCAPS assays are listed in Tables S1-S3. Post PCR, we digested the amplicons either with *Nsi*I (*ddm1-1*) or *Rsa*I (*ddm1-2*) restriction enzyme according to conditions suggested by *New England Biolabs*. Both *ddm1-1* and *ddm1-2* carry G to A base substitutions, resulting in the loss of *Nsi*I and *RsaI* sites, respectively. Therefore, PCR amplicons from these mutants are not digested in the presence of *Rsa*I or *Nsi*I enzymes, while those from WT-Col-0 are digested by these enzymes. The digested products were resolved using 3% agarose gel electrophoresis. For genotyping *fas1-4* (SAIL_662_D10), *fas2-4* 9 (SALK_012874), and *cmt2-3* (SALK_033228) mutants, which carry T-DNA insertions, we used PCR assays to amplify either a WT product or T-DNA using primers, synthesized according to the SALK T-DNA Primer Design Tool (http://signal.salk.edu/tdnaprimers.2.html), and the sequences of the primers used are listed in Table S1.

### Cloning and sequencing of *ddm1-2* RNA

Primers designed by a previous study were used to amplify the Exon 11-flanking region in *ddm1-2* mutants (25). The PCR product was gel-purified using a PCR clean-up kit (Macherey-Nagel; Cat. No. 740609.50). The purified PCR Product was cloned into pGEM^®^-T easy vector (Promega Cat. No. A1360) and was sequenced by Sanger sequencing to confirm the splicing defect in *ddm1-2* mutants, resulting in premature translation termination of DDM1 protein.

### RT PCR

For RT-PCR, four-week-old leaf tissues were used to extract RNA using a *Qiagen RNeasy plant mini kit* (Cat. No. 74904). The isolated RNA was treated with Invitrogen’s TURBO DNA-free™ kit (Cat. No. AM1907) to degrade contaminating genomic DNA. For cDNA synthesis with random hexamers, Invitrogen’s SuperScript™ III first-strand synthesis kit (Cat. No. 18080051) was used, starting with 1000 ng of DNA-free total RNA in a 20 μl reaction. One microliter of cDNA from this reverse transcription reaction was then used for PCR amplification of rRNA subtypes and ACTIN 2/7 control.

### Primers, PCR profiles, and agarose gel electrophoresis of PCR products

PCR amplification of rRNA variants, NOR-linked markers, and any other amplicons used in this manuscript was carried out using primers listed in Table S1 and PCR profiles listed in Table S2. Enzymes used in CAPS and dCAPS assays are listed in Table S3. PCR and RT-PCR products were resolved by electrophoresis on 1%–2.5% agarose gels in TAE buffer.

### Preparation of agarose gel image panels

Agarose gel images were captured and processed using Image Lab software (Bio-Rad) and compiled using Microsoft Office PowerPoint tool. Wherever data from a subset of representative plant samples were shown, gel images from the respective samples have been spliced to create image panels.

## Results

### PCR-based assays to determine the occurrence and the extent of rDNA genomic instability

In the Col-0 ecotype, rRNA genes at NORs are arranged as tandem repeats in head-to-tail fashion facing the centromere (Fig. 1a) (7). PCR amplification of length-variable 3′ ETS of rRNA genes yields four distinct variants, named VAR1-4 (Fig. 1b, first lane) (8, 10). A recent study in the Col-0 ecotype identified 74 rRNA gene subtypes, based on length and sequence variation among the 45S rRNA genes (7). Nonetheless, the primers that amplify the four rRNA subtypes bind to most of the repeats from both NORs. Of these, VAR1 constitutes the largest fraction, and most of them are located on *NOR2*, which undergoes selecting silencing during development (Fig. 1b, second lane) (8–10). The VAR2 genes map primarily to *NOR4,* and a subset of VAR3 genes is present on *NOR2,* while the rest along with a small fraction of VAR4 genes are located on *NOR4*, which remains transcriptionally active during development (8, 9). Primarily we use VAR1 and VAR2 genes as *NOR2* and *NOR4* representatives, respectively, in genetic mapping studies

**Fig. 1.**
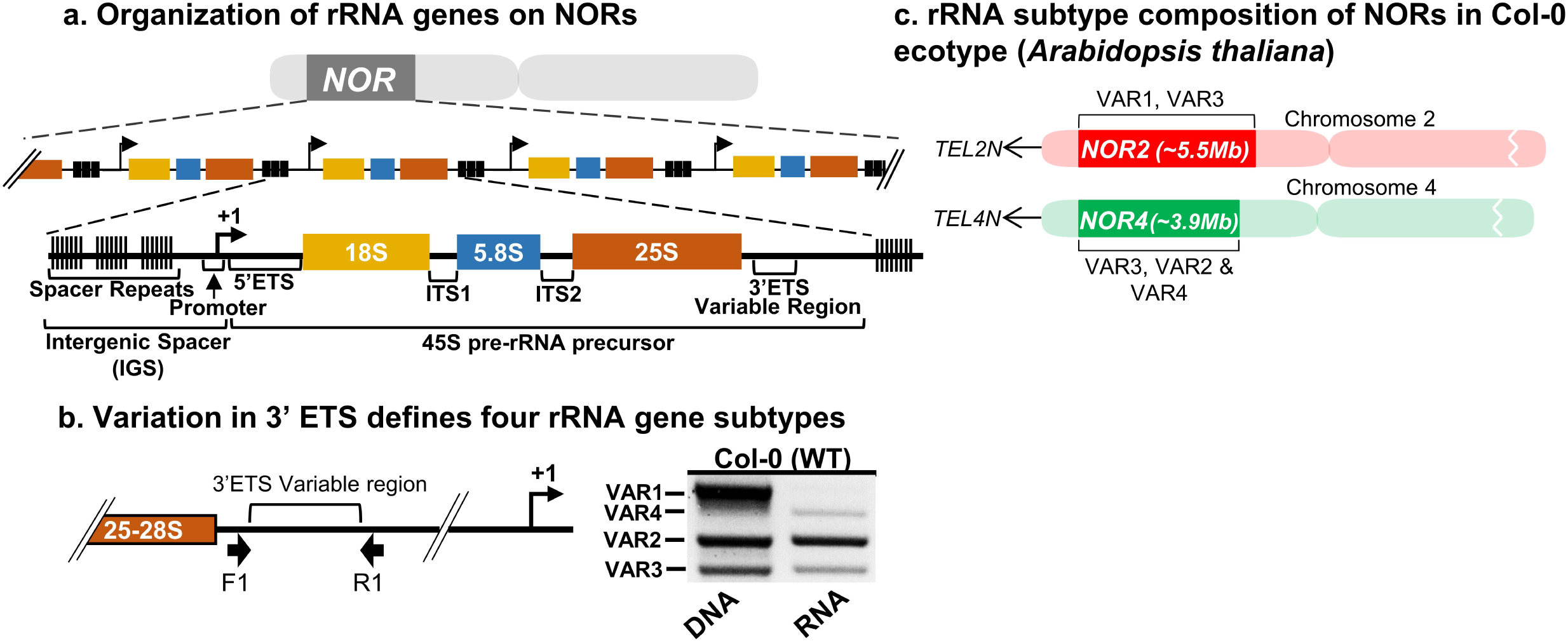
45S rRNA gene organization, rRNA subtypes and their localization in *Arabidopsis thaliana.* **a** the arrangement of rRNA genes as tandem repeats on NORs. **b** Variation in the 3’ETS region of rRNA genes defines four subtypes in the ecotype Col-0 of *Arabidopsis thaliana.* The gel images on the right side show PCR amplification of rRNA variants from genomic DNA and cDNA. **c** Diagram depicts the localization of rRNA gene subtypes on *NOR2* and *NOR4* of A *thaliana* (Col-0).

In Col-0, both the NORs are capped by telomeres, and unique sequences exist in the NOR-telomere junctions. Using this polymorphism, the NOR2– and NOR4-telomere junctions, named *NOR2-TEL2N* and *NOR4-TEL4N*, respectively, can be uniquely amplified using multiplex PCR (Fig. S1a-b) (9). In Col-0 *NOR4*, 3^rd^ and 25^th^ gene repeats carry large deletions (7). We designed primers to uniquely amplify these regions as Col-0 NOR4-specific markers, labelled N4-m1 and N4-m2, respectively (Fig. S1 c-d). Of these, N4-m2 is ∼0.25 Mb away from *NOR4-TEL4N* (Fig. S1c). In *NOR2*, the second and the third gene repeats are inverted (7), resulting in distinct sequences between the first and the second repeat, and the third and the fourth repeat, when compared to the other tandem rDNA repeats (Fig. S1e). Similarly, we designed primers to uniquely amplify these junction sequences as Col-0 NOR2-specific markers, labelled N2-m1 and N2-m2, respectively (Fig. S1f). Of these, N2-m2 is ∼35 kb away from *NOR2-TEL2N* (Fig. S1e). In Arabidopsis mutants, first, we used PCR assays described in Fig. 1c and Fig. S1 to detect rDNA genomic instabilities, if any. Upon detecting noticeable rDNA instability, we carried out genetic mapping of the rRNA variants and the NOR-telomere junctions to further characterize the instability patterns.

### The *ddm1-2* mutants display loss of ∼ 0.25 Mb of *NOR4* genes and the associated telomere in a stochastic manner

The *ddm1-2* mutants carry a G to A substitution, resulting in a defective splicing that led to the synthesis of a truncated protein due to the introduction of a premature stop codon in the reading frame of the gene (25) (Fig. S2). Initially, we amplified NOR-telomere junctions in ∼50 individual *ddm1-2* mutant plants using their genomic DNA through multiplex PCR (Fig. S1a-b) (9). Of these, some plants showed loss of *NOR4-TEL4N* (Fig. 2, topmost gel images, plant no. 5, 14, 16, 37, 42, and 50), and we confirmed these plants to be true *ddm1-2* mutants through dCAPS-based genotyping (Fig. S3) (36). Next, we sought to determine if *NOR4* genes have been deleted in the *ddm1-2* mutants that showed loss of the *NOR4-TEL4N* (from hereafter, those mutants will be referred to as *ddm1-2(ΔNOR4-TEL4N)*. We tried PCR amplifying the NOR4-specific markers N4-m1 and N4-m2 in these mutants. The *ddm1-2(ΔNOR4-TEL4N)* plants failed to amplify N4-m1 and N4-m2 markers (Figs. 2 and S5, 2^nd^ and 3^rd^ panels of gel images), indicating that these mutants have lost at least ∼ 0.25 Mb of telomere-proximal *NOR4* rDNA. Further, PCR amplification of rRNA variants in the same set of mutant plants revealed some reduction in VAR3 genes in them when compared to other plants (Fig. 2, bottom most panel of gel images, plant no. 5, 14, 16, 37, 42, and 50). This reduction VAR3 content is due to the fact that a stretch of VAR3 genes are present adjoining the *NOR4* telomere (7). PCR amplification of some centromere-proximal regions, a few kb away from *NOR4* (Fig. S4), suggests that the rDNA loss observed in *ddm1-2(ΔNOR4-TEL4N)* mutants is likely restricted to the *NOR4* region. Next, in another 110 *ddm1-2* plants in the successive generation (G_n+1_), we PCR-amplified NOR-telomere junctions to determine whether the loss of the other NOR-telomere junction (*NOR2-TEL2N)* has occurred as well. In this pool of plants also, we found only the loss of *NOR4-TEL4N* (Fig. S6).

**Fig. 2.**
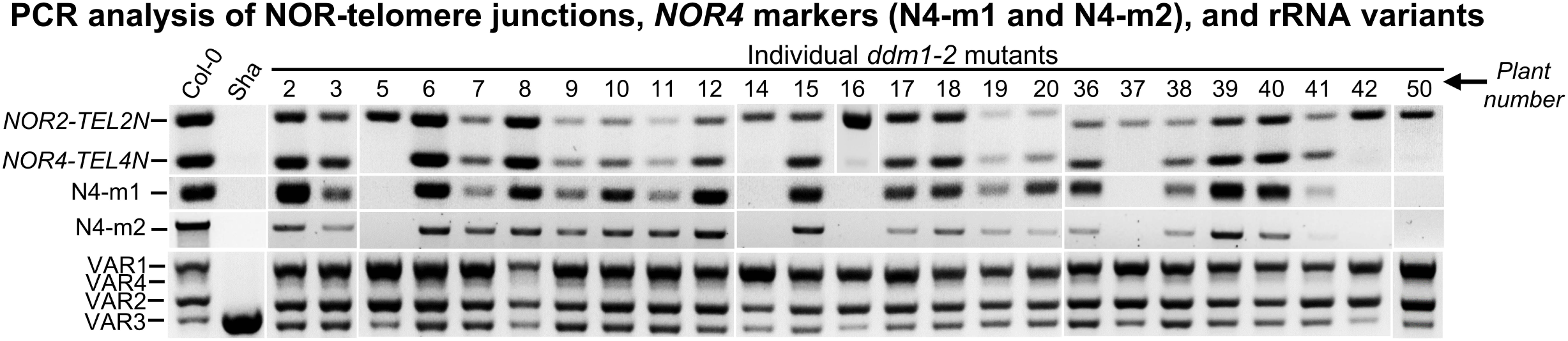
The *ddm1-2* mutants display genomic instability of 45S rRNA genes and its associated telomeres in a stochastic manner. The gel images show PCR analysis of NOR-telomere junctions, *NOR4* markers (N4-m1 and N4-m2), and rRNA gene subtypes, in WT-Col-0, Sha, and several individual *ddm1-2* mutant plants. About 50 individual *ddm1-2* mutant plants were screened, of which data are shown for a subset, while the data for remaining plants are shown in Fig. S5.

### The *ddm1-2(ΔNOR4-TEL4N)* mutants show *NOR4* conversion involving translocation of *NOR2* genes to *NOR4*

Next, we hypothesized that the lost *NOR4* rDNA in *ddm1-2(ΔNOR4-TEL4N)* mutants could have been repaired using the corresponding *NOR2* sequences, similar to what was observed in Arabidopsis *atxr5 atxr6* mutants (9). To test this hypothesis, we crossed the *ddm1-2(ΔNOR4-TEL4N)* mutants with ecotype Shahdara (Sha), which contains exclusively VAR3 subtype rRNA genes, and their NOR-telomere junction sequences are different when compared to those of Col-0 sequences. Therefore, genetic mapping of VAR1, VAR2, and NOR-telomere junctions is possible in a mapping population derived from this cross. The resulting F_1_ hybrids from the Sha x *ddm1-2(ΔNOR4-TEL4N)* cross were then allowed to self-fertilize to obtain an F_2_ progeny. About 100 F_2_ plants were grown, and genomic DNA (gDNA) was isolated from ∼4-week-old plants and PCR-amplified for rRNA gene variant content, the NOR-telomere junctions, and the NOR-adjacent markers that discriminate Col-0 from Sha chromosomes. The mapping strategy outlined in Figure 3a had been used before (8, 9). However, we employed recently-designed NOR-linked markers, which exploited the length-based polymorphism in the NOR-flanking regions of Col-0 and Sha ecotypes (Fig. 3b) (37).

**Fig. 3.**
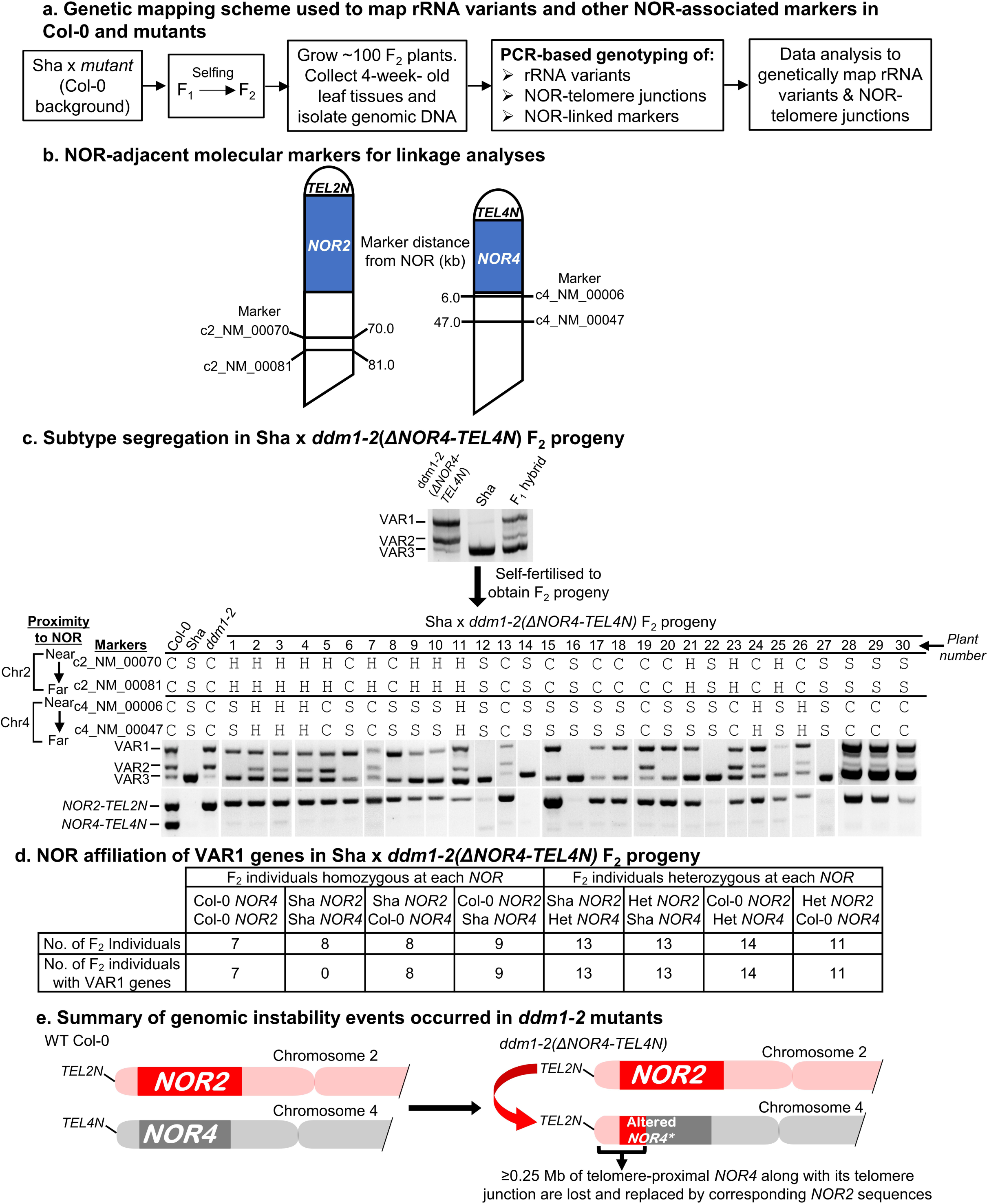
In *ddm1-2* mutants, VAR1 genes and *NOR2-TEL2N* map to *NOR2* and *NOR4* instead of *NOR2* only. **a** The flow chart shows the genetic scheme used to map the rRNA variants and the NOR-associated markers. **b** Positions of telomeres, NORs, and genetic markers on the northern ends of chromosomes 2 and 4. Marker names and distances (in kb) from the NORs are shown. **c** The top gel image shows rRNA variant PCR of the parent plants (Sha and *ddm1-2(./J.NOR4-TEL4N)* and the F_1_ plant used for selfing. The panels below show segregation analysis of Col-0 rRNA gene subtypes, NOR-telomere junctions, and ecotype-specific NOR-adjacent molecular markers among F2 progeny of Sha x *ddm1-2(./J.NOR4-TEL4N.* Segregation data for the molecular markers is provided in Fig. S7. **d** Summary of Sha x *ddm1-2(./J.NOR4-TEL4N)* F2 progeny segregation data for informative genotypes. **e** Graphical summary of genomic instability of NORs (loss and replacement) occurred in *ddm1-2* mutants compared to WT Col-0 NORs. C=Col-0; S=Sha; H=heterozygous.

From the previous studies, it was established that the VAR1 genes map mainly to *NOR2* in the wild-type (WT) Col-0 (8, 9), but in F_2_ progeny of Sha x *ddm1-2(ΔNOR4-TEL4N)* cross, we found that VAR1 genes map to both *NOR2* and *NOR4*, as indicated by their cosegregation with both NOR2– and NOR4-linked markers. For example, in Figures 3c and S7, F_2_ plants numbered 17 and 18 are homozygous for Col-0 NOR2-linked markers and Sha NOR4-linked markers, indicating that they carry Col-0 *NOR2* and Sha *NOR4*. As expected, these plants show the presence of VAR1 genes and *NOR2-TEL2N* on *NOR2* (Fig. 3c). However, F_2_ plants that are homozygous for Sha *NOR2* and Col-0 *NOR4* (Fig. 3c-d and Fig. S7, e.g., F_2_ plants no. 29 and 30) also showed the presence of VAR1 genes and Col-0 *NOR2-TEL2N* (Fig. 3c). Collectively, these results establish that VAR1 genes and *NOR2-TEL2N* are located both on *NOR2* and *NOR4* in *ddm1-2(ΔNOR4-TEL4N)* plants (Fig. 3c-d) unlike the WT Col-0 plants where VAR1 genes are primarily NOR2-localized. The results demonstrate the occurrence of a translocation of *NOR2* VAR1 genes and the associated telomere *NOR2-TEL2N* to NOR4 in *ddm1-2(ΔNOR4-TEL4N)* plants. A diagram graphically illustrating genomic instability of *NOR4* and *NOR4-TEL4N* (loss and replacement) occurred in *ddm1-2(ΔNOR4-TEL4N)* mutants compared to WT Col-0 NORs, is shown in Fig. 3e.

Further, we sought to determine the extent of *NOR2* rDNA translocated to *NOR4*, through PCR amplification of NOR2-specific markers N2-m1 and N2-m2 in the informative F_2_ plants of Sha x *ddm1-2(ΔNOR4-TEL4N)* and the Sha x WT Col-0 crosses (control). First, we show that N2-m1 and N2-m2 markers are present in the informative F_2_ plants of Sha x WT-Col-0 cross, which carry Col-0 *NOR2* and Sha *NOR4* (Fig. S8, plants no. 1 and 2), but not in the F_2_ plants that carry Sha *NOR2* and Col-0 *NOR4* (Fig. S8, plants numbered 3 and 4). These results are as expected. However, F_2_ plants of Sha x *ddm1-2(ΔNOR4-TEL4N)* cross showed the presence of Col-0 *NOR2*-specific markers N2-m1 and N2-m2, irrespective of whether they are homozygous for Col-0 *NOR2* or *NOR4* (Fig. S8, plants no. 11, 12, 13, and 15). Note that N2-m1 and N2-m2 markers are Col-0 specific (absent in Sha ecotype). So, the results indicate that at least ∼35 kb of *NOR2* rDNA, along with *NOR2-TEL2N,* has been translocated to *NOR4* in *ddm1-2(ΔNOR4-TEL4N)* plants, replacing ∼0.25 Mb of lost telomere-proximal *NOR4* rDNA and *NOR4-TEL4N*. A chi-square test for genotyping data of a NOR2– and a NOR4-linked marker showed that the markers are segregating in a Mendelian fashion (Fig. S9), indicating that meiotic segregation of the homologous chromosomes in the F_2_ progeny of Sha x *ddm1-2(ΔNOR4-TEL4N)* is normal.

### *ddm1-1* mutants did not show any observable rDNA genomic instability

The *ddm1-1* mutants carry a G to A base substitution in exon 13, resulting in a C615Y (cysteine to tyrosine) missense mutation in the Helicase Superfamily C terminal (HELICc) domain of DDM1 protein (Fig. S10a) (23). WT Col-0 plants can be distinguished from the *ddm1-1* mutants by a CAPS assay (Fig. S10b-c). To detect rDNA genomic instability, if any, we PCR-amplified NOR-telomere junctions in ∼100 *ddm1-1* plants, but we could not detect any loss of NOR-telomere junctions in them (Fig. S11a). Then, we propagated these mutants for three more generations and again tested ∼100 mutant plants for rDNA genomic instability through PCR amplification of NOR-telomere junctions and rRNA variants. Yet we could not detect any observable rDNA genomic instability in these mutants (Fig. S11b).

### The rDNA instability patterns influence the rRNA variant expression patterns in *ddm1-2* mutants

To test the effects of rDNA instability in *ddm1-2* mutants on rRNA variant expression pattern, we carried out RT-PCR analysis of rRNA variants and actin (as an internal control) in *ddm1-2* mutants with no loss of NOR-telomere junctions (from hereafter referred to as *ddm1-2(no NT loss)* and *ddm1-2(ΔNOR4-TEL4N)* plants, along with WT Col-0 controls. As expected, the VAR1 (*NOR2*) genes were silenced in WT Col-0 (Fig. 4a, lane 1). By contrast, we observed significant expression of VAR1 (*NOR2*) genes in both groups of *ddm1-2* mutants (Fig. 4a, lanes 2-7). However, comparison of rRNA expression patterns between *ddm1-2(no NT loss)* and *ddm1-2(ΔNOR4-TEL4N)* mutant plants revealed two differences. First, relatively more VAR1 gene expression was seen in *ddm1-2(ΔNOR4-TEL4N)* compared to *ddm1-2 (no NT loss)* (Fig. 4a, compare lanes 2-4 with lanes 5-7), reflecting the effect of translocation of some *NOR2* VAR1 genes to *NOR4* in *ddm1-2(ΔNOR4-TEL4N)* plants on VAR1 rRNA levels. Second, VAR3 expression levels were lower in *ddm1-2(ΔNOR4-TEL4N)* mutant plants than *ddm1-2 (no NT loss)* (Fig. 4a, compare lanes 2-4 with lanes 5-7), reflecting some loss of telomere-proximal *NOR4* VAR3 genes in *ddm1-2(ΔNOR4-TEL4N)*.

**Fig. 4.**
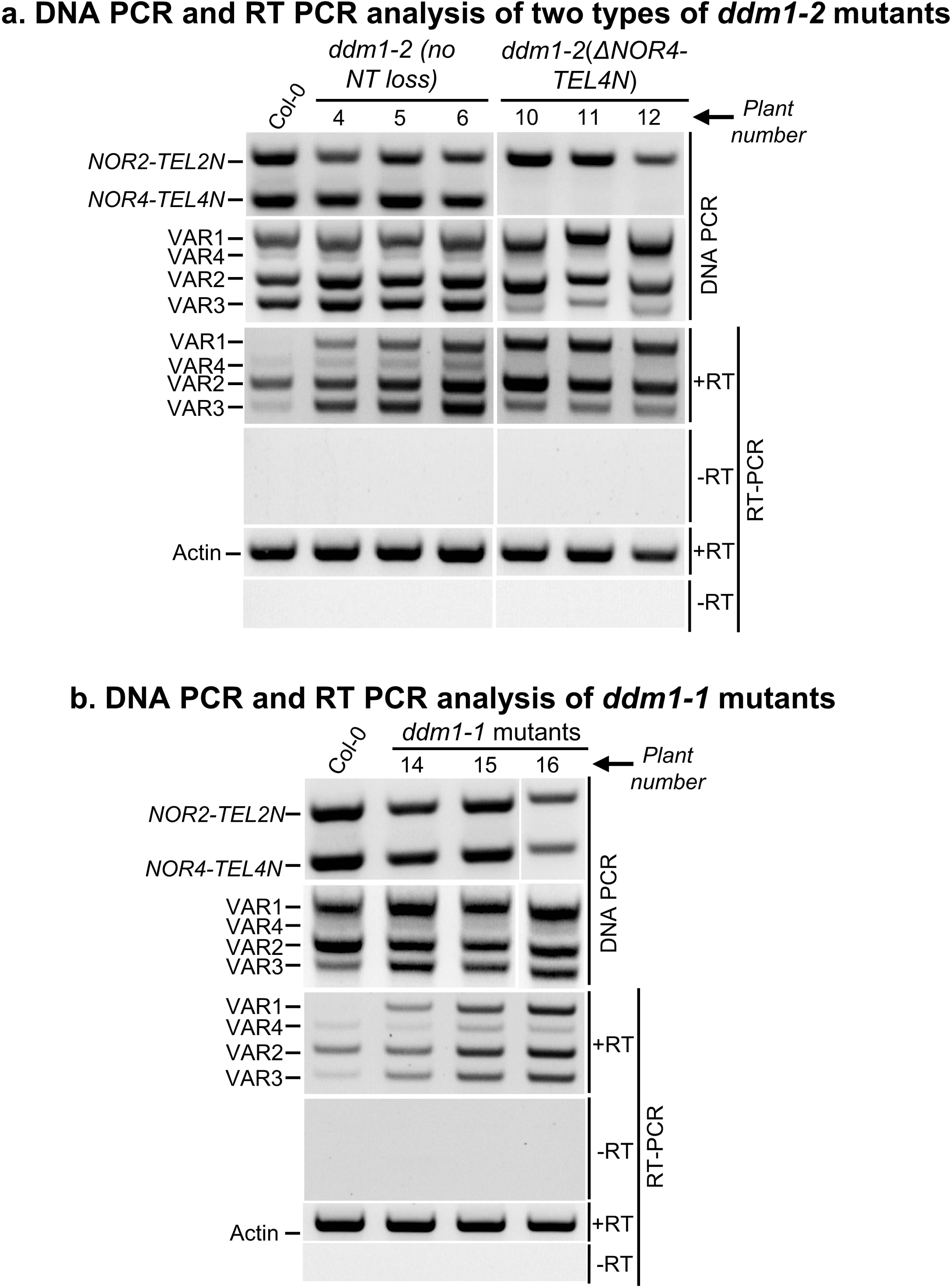
rRNA variant expression pattern in *ddm1-2* mutants reflects the rDNA instability pattern and *ddm1-1* mutants show release of rRNA gene silencing without any observable rDNA instability. Panels of gel images show PCR analysis of gDNA for NOR-telomere junctions and rRNA variants, and RT-PCR analysis of rRNA variants and Actin (as a control) in WT Col-0 and **(a)** *ddm1-2* mutants or (**b**) *ddm1-1* mutants.

### Disruption of selective *NOR2* gene silencing was observed in *ddm1-1* mutants

Similar RT-PCR analysis of rRNA variants in *ddm1-1* mutants revealed significant VAR1 expression (disruption of *NOR2* gene silencing) in *ddm1-1* mutants also (Fig. 4b), despite the absence of any observable rDNA genomic instability in them (Fig. S11). These results underscore the importance of the HELICc domain in DDM1 protein in selective rRNA silencing (Fig. S12a-b).

The *ddm1-2(ΔNOR4-TEL4N)* plants displayed rDNA genomic instability in some plants and correspondingly altered rRNA variant expression pattern. While the *ddm1-2 (no NT loss)* plants displayed *NOR2* gene silencing disruption with no apparent rDNA instability. Of these two mutant phenotypes, *ddm1-1* mutants displayed only the latter. These differential mutant phenotypes can be attributed to differences among the mutant alleles. The *ddm1-2* carries a truncated DDM1 protein with a 240 amino acid (aa) deletion and an 18 aa insertion, followed by a stop codon, in the C-terminal domain, resulting in the loss of at least two functional domains; a conserved histone 2A variant W (H2A.W) binding site and the HELICc domain. (Fig. S12 a&c) (23, 24). On the other hand, *ddm1-1* carries just one aa substitution (C615Y) in the HELICc domain, that is encoded by exon 12-14 sequences (Fig. S12b). This one aa substitution (tyrosine replacing cysteine) abolishes the function of the HELICc domain by disrupting the disulfide formation, a key requirement for its function (23). This mutation is also known to affect DNA methylation levels in all cytosine contexts (25), further underscoring the importance of C615 in the HELICc domain in DDM1 functionality.

### Loss of CAF-1 leads to loss of *NOR2* and *NOR4* genes and the associated telomeres in a stochastic manner

Previous studies have shown that the loss of CAF-1 function leads to loss of rDNA and release of *NOR2* gene silencing (33, 34). However, it remains to be determined whether the observed rDNA instability involves any translocation of rDNA and the associated telomeres from one NOR to the other, and if any, whether such rDNA translocations would influence rRNA gene expression patterns. To address this question, we studied two *caf-1* mutants, *fas1-4* and *fas2-4*, which carry T-DNA insertions in *FAS1* and *FAS2* genes, respectively (Fig. S13). *FAS1* and *FAS2* constitute members of the CAF-1 heterotrimer complex (27). PCR amplification of NOR-telomere junctions in ∼100 plants each in *fas1-4* and *fas2-4* mutants revealed a loss of *NOR2-TEL2N* (Fig. 5a, plant# 5 and 21; Fig. 5b, plant#3, 4, 9, 10, 12, and 15) or *NOR4-TEL4N* in a stochastic manner (Fig. 5a, plant# 9 and 20; Fig. 5b, plant# 18, 19, 22, and 25). From hereafter, *fas1-4* and *fas2-4* mutants, which have lost *NOR2-TEL2N,* will be referred to as *fas1-4*(*ΔNOR2-TEL2N*) and *fas2-4*(*ΔNOR2-TEL2N*), respectively. Similarly, those mutant plants that showed the loss of *NOR4-TEL4N* will be referred to as *fas1-4*(*ΔNOR4-TEL4N*) and *fas2-4*(*ΔNOR4-TEL4N*), respectively. In some mutants, we did not detect any loss of NOR-telomere junctions (hereafter referred to as *fas1-4(no NT loss)* and *fas2-4(no NT loss)*), but we did observe a reduction in some rRNA variant content. For example, in *fas1-4*, plant# 16, 2, 17, and 7 showed reduced VAR2 content when compared to WT-Col-0 plants (Fig. 5a, bottom-most panel of gel images. Compare relative band intensities of VAR2 and VAR1 genes between WT-Col-0 and the mutants). Similar results were obtained in *fas2-4* mutants. (Fig. 5b, bottom-most panel of gel images, plant# 2, 17, and 26. Compare relative band intensities of VAR2 and VAR1 genes between WT Col-0 and the mutants).

**Fig. 5.**
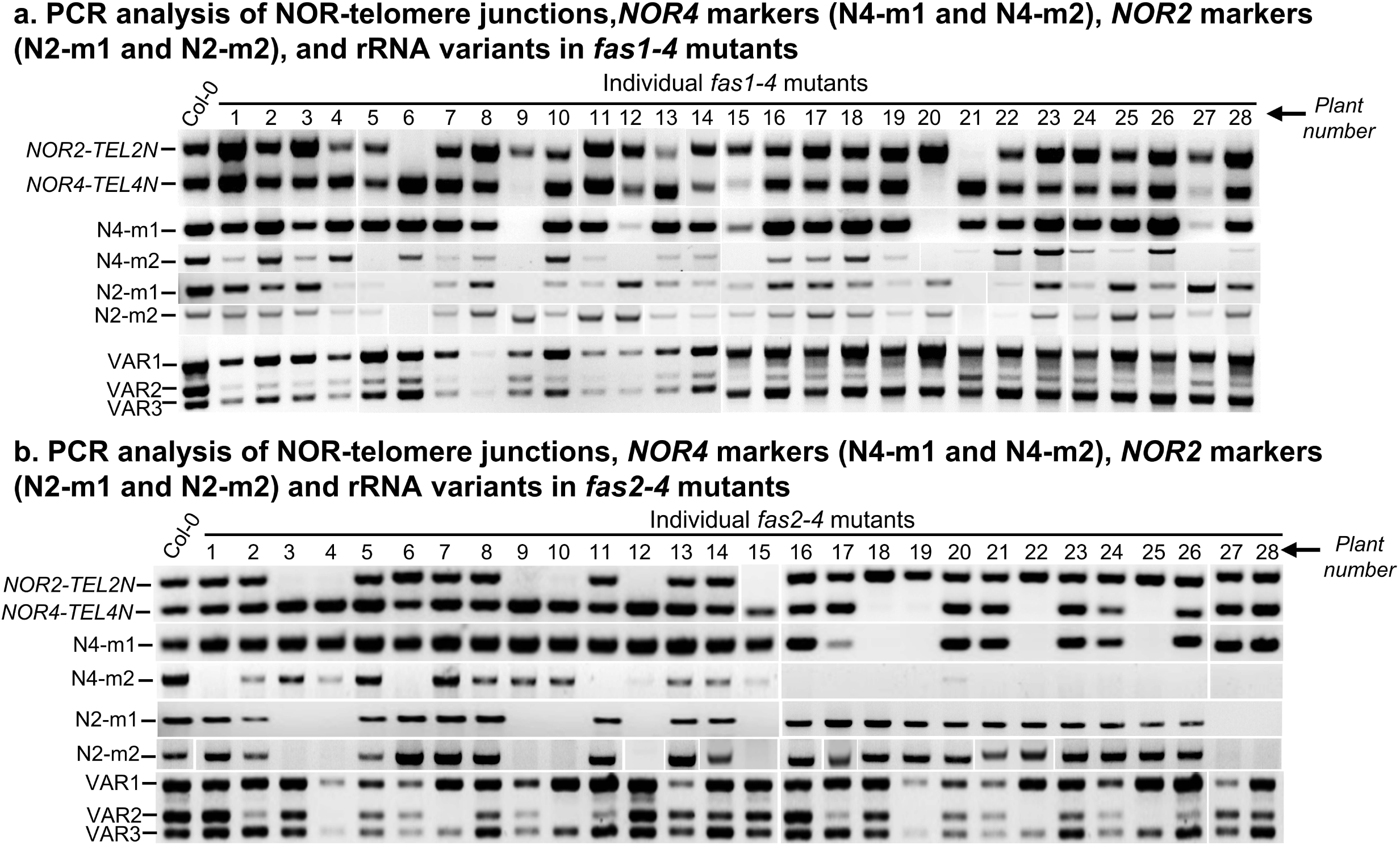
The *caf-1* mutants display genomic instability of 45S rRNA genes and the associated telomeres in a stochastic manner. Panels of gel images show PCR analysis of NOR-telomere junctions, *NOR4* markers (N4-m1 and N4-m2), NOR2 markers (N2-m1 and N2-m2), and rRNA variants in WT-Col-0 and (**a**) fast *1-4* mutants or (**b**) fast *2-4* mutants.

Next, using PCR amplification of the *NOR2* markers N2-m1 and N2-m2, we show that the *fas1-4*(*ΔNOR2-TEL2N*) and *fas2-4*(*ΔNOR2-TEL2N*) mutants have lost at least 35 kb of telomere-proximal rDNA along with the telomere *NOR2-TEL2N* (Fig. 5a, plant# 6 and 21; Fig. 5b, plant#3, 4, 9, 10, 12, and 15). Similarly, through PCR amplification of the *NOR4* markers N4-m1 and N4-m2, we show that the *fas1-4*(*ΔNOR4-TEL4N*) and *fas2-4*(*ΔNOR4-TEL4N*) mutants have lost at least ∼0.25 Mb of telomere-proximal rDNA along with the telomere *NOR4-TEL4N* (Fig. 5a, plant# 9 and 20; Fig. 5b, plant# 18, 19, 22, and 25). Owing to these phenomena, the mutants displayed altered rRNA gene variant profile at the DNA level, reflecting varying degrees of genomic instability and its stochastic nature, irrespective of the loss of NOR-telomere junctions (Fig. 5a-b, bottom-most panel of gel images). PCR amplification of a few markers flanking the NORs indicated that the genomic instability is likely restricted to the NOR regions in the *fas* mutants (Fig. S14).

### The *caf-1* mutants display NOR conversion involving the translocation of rRNA genes and the associated telomeres from one NOR to the other

The *caf-1* mutants displayed stochastic loss of both *NOR2-TEL2N* and *NOR4-TEL4* along with the adjoining rDNA. To test if these genomic instabilities involved exchanges of rDNA between NORs, we crossed both *fas1-4(ΔNOR4-TEL4N)* and *fas1-4(ΔNOR2-TEL2N)* plants with Sha ecotype. The resulting F_1_ hybrids were allowed to self-fertilize to obtain F_2_ progeny. ∼ 100 F_2_ plants were grown, and when they were ∼4-week-old, tissues were harvested to isolate gDNA, which was then tested for rRNA gene variant content, the NOR-telomere junctions, and the NOR-adjacent markers that discriminate Col-0 from Sha chromosomes. The mapping strategy outlined in Figure 3a was used for mapping the rRNA variants and the NOR-telomere junctions in the F_2_ progenies of both *fas1-4(ΔNOR4-TEL4N)* x Sha and *fas1-4(ΔNOR2-TEL2N)* x Sha crosses. The NOR-linked markers used for mapping are given in Figure 6a.

**Fig. 6.**
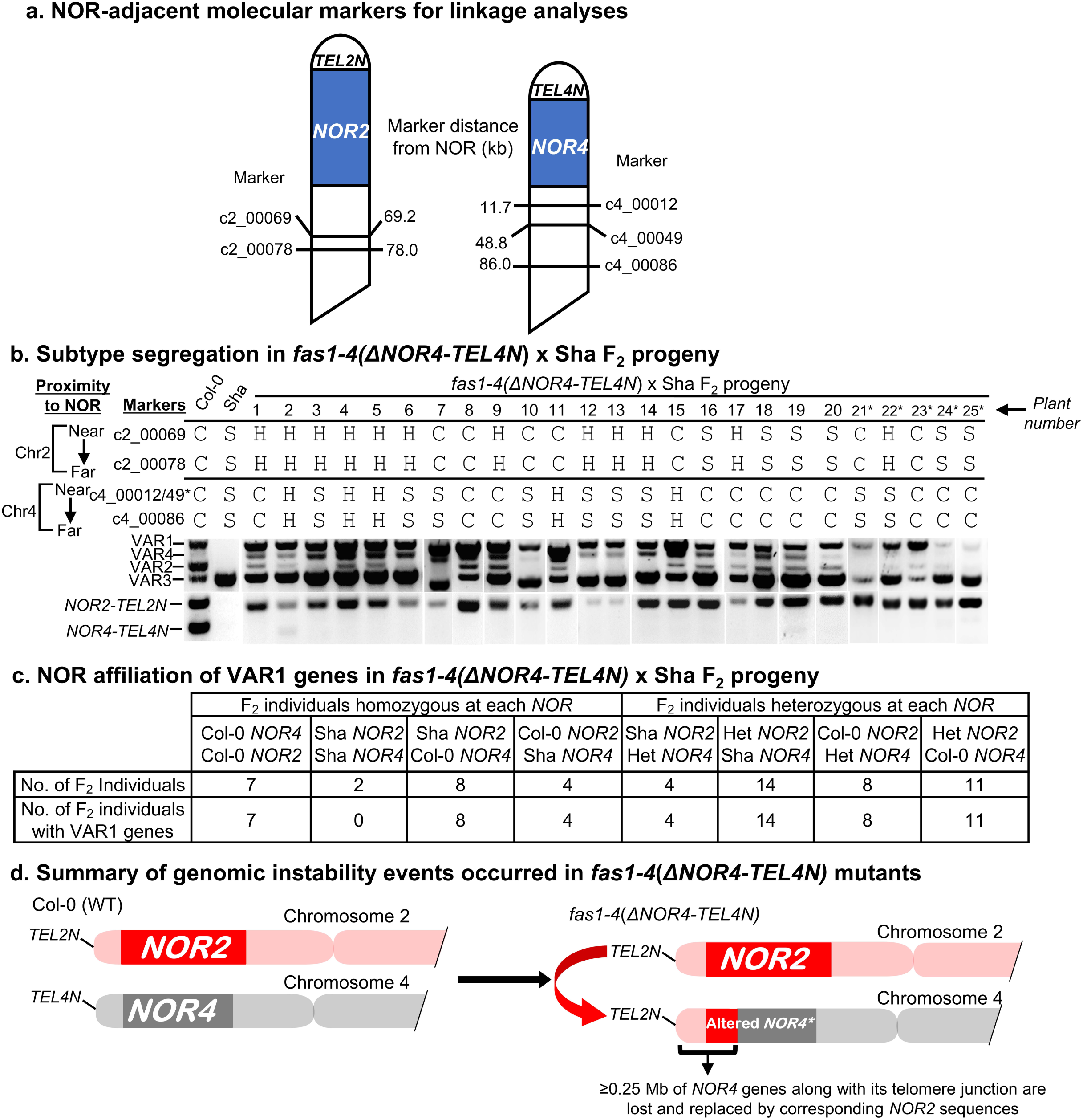
In *fas1-4(Δ.NOR4-TEL4N)* mutants, VAR1 genes and *NOR2-TEL2N* map to *NOR2* and *NOR4* instead of *NOR2* only. **a** Positions of telomeres, NORs, and genetic markers on the northern ends of chromosomes 2 and 4. Marker names and distances (in kb) from the NORs are shown. **b** The panel shows segregation analysis of Col-0 rRNA gene subtypes, NOR-telomere junctions, and ecotype specific NOR-adjacent molecular markers among F_2_ progeny of *fas1-4(iJ.NOR4-TEL4N)* x Sha. Segregation data for the molecular markers is provided in Fig S15. **c** Summary of *fas1-4(iJ.NOR4-TEL4N)* x Sha F_2_ progeny segregation data for informative genotypes. **d** Graphical summary of genomic instability of NORs (loss and replacement) occurred in *fas1-4(iJ.NOR4-TEL4N)* mutants compared to WT-Col-0 NORs. C=Col-0; S=Sha; H=heterozygous.

In WT Col-0 plants, VAR1 genes mainly map to *NOR2* (8, 9), but in F_2_ progeny of *fas1-4(ΔNOR4-TEL4N)* x Sha cross, we found that VAR1 genes map to both *NOR2* and *NOR4*, as indicated by their cosegregation with both NOR2– and NOR4-linked markers. For example, in Figures 6 and S15, F_2_ plants #7, 10, and 21 are homozygous for Col-0 NOR2-linked markers and Sha NOR4-linked markers, indicating that they carry Col-0 *NOR2* and Sha *NOR4*. As expected, these plants show the presence of VAR1 genes on *NOR2* and Col-0 *NOR2-TEL2N*. However, F_2_ plants that are homozygous for Sha *NOR2* and Col-0 *NOR4* (Fig. 6b and Fig. S15, e.g., F_2_ plants# 16, 20, and 21) also show the presence of VAR1 genes and Col-0 *NOR2-TEL2N*. Collectively, these results establish that the VAR1 genes and the *NOR2-TEL2N* are located both on *NOR2* and *NOR4* in *fas1-4(ΔNOR4-TEL4N)* plants (Fig. 6c-d), indicating a translocation of *NOR2* VAR1 genes and the associated telomere *NOR2-TEL2N* to NOR4 in *fas1-4(ΔNOR4-TEL4N)* plants (Fig. 5a, topmost panel of gel images).

Next, we tested for the extent of *NOR2* rDNA translocated to *NOR4*, through PCR amplification of *NOR2* markers N2-m1 and N2-m2. First, we show that N2-m1 and N2-m2 are present in F_2_ plants derived from the Sha x WT Col-0 cross, which carry Col-0 *NOR2* and Sha *NOR4* (Fig. S16, plant no. 1 and 2), but not in the F_2_ plants which carry Sha *NOR2* and Col-0 *NOR4* (Fig. S16, plant no. 3 and 4), as expected. However, F_2_ plants of *fas1-4(ΔNOR4-TEL4N)* x Sha showed the presence of N2-m1 and N2-m2 markers whether they carry Col-0 *NOR2* or *NOR4* (Fig. S16, plant no. 12, 13, 21, 25, and 24), indicating that at least ∼35 kb of *NOR2* rDNA along with *NOR2-TEL2N* has been translocated to *NOR4* in *fas1-4(ΔNOR4-TEL4N)* plants. A chi-square test for genotyping data of a NOR2– and a NOR4-linked marker showed that the markers are segregating in Mendelian fashion (Fig. S17), indicating that meiotic homologous chromosome segregation of in F_2_ plants of *fas1-4(ΔNOR4-TEL4N)* x Sha has been normal.

Similarly, mapping of rRNA variants and the NOR-telomere junctions was carried out in F_2_ progeny of *fas1-4(ΔNOR2-TEL2N)* x Sha. The results showed similar loss of *NOR2* rDNA and *NOR2-TEL2N*, and their replacement with the corresponding *NOR4* rDNA and the associated telomere (Figs. 7 and S18-S19) sequences. Chi-square test results in the same F_2_ progeny indicated that meiotic segregation has been normal in them.

**Fig. 7.**
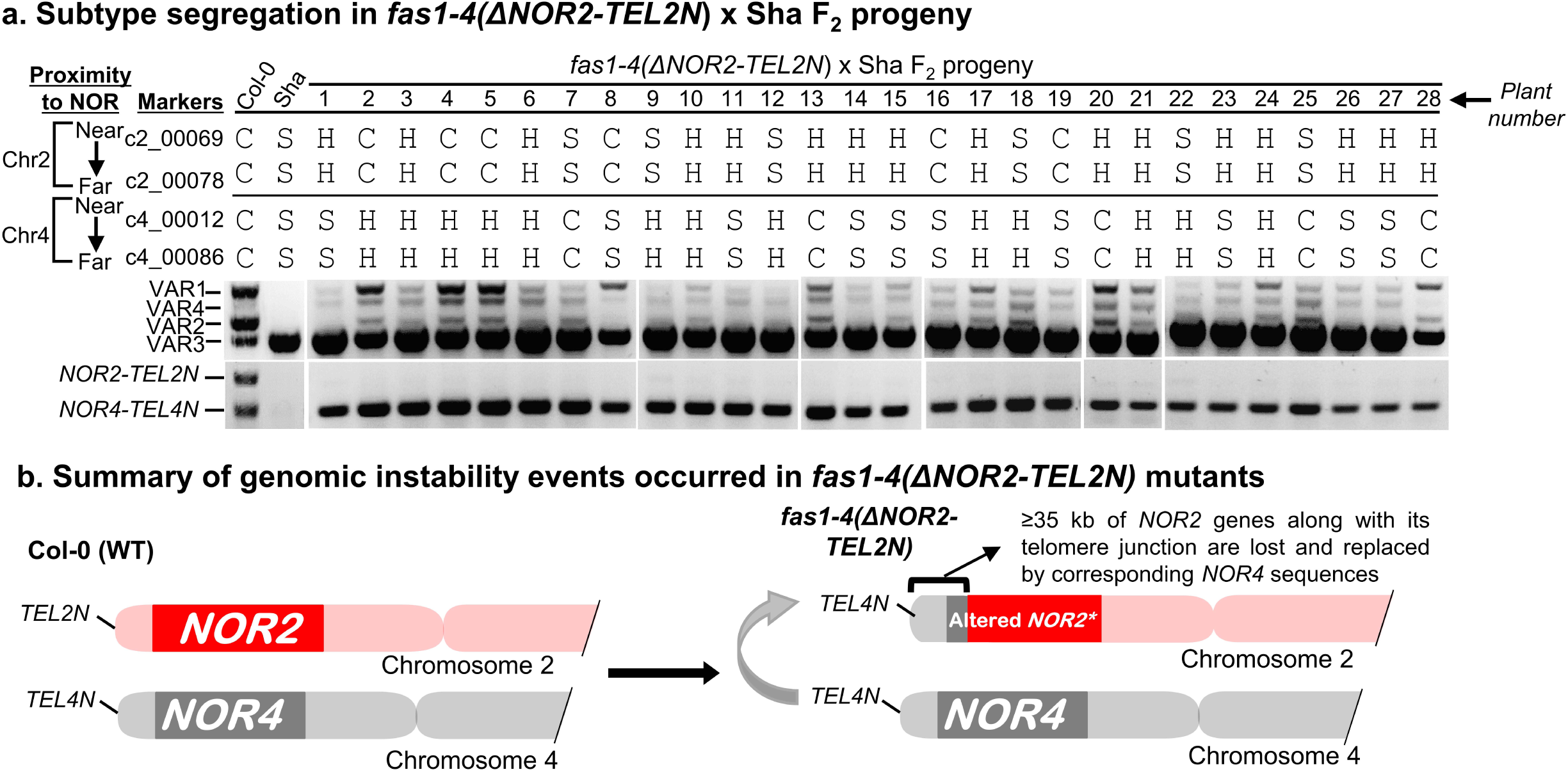
In *fas1-4(lJ.NOR2-TEL2N)* mutants, *NOR4-TEL4N* map to *NOR2* and *NOR4* instead of *NOR4* only. **a** The panels show segregation analysis of Col-0 rRNA gene subtypes, NOR-telomere junctions, and ecotype-specific NOR-adjacent molecular markers among **F**_2_ progeny of *fas1-4(lJ.NOR2-TEL2N)* x Sha. Segregation data for the molecular markers is provided in Fig S18. **b** Graphical summary of genomic instability of NORs (loss and replacement) occurred in *fas1-4(lJ.NOR2-TEL2N)* mutants compared to WT-Col-0 NORs.

Based on the occurrence of genomic instabilities involving rDNA and the associated telomeres in *fas2-4* mutants (Fig. 5b), we presume that NOR conversion events akin to those observed in *fas1-4* mutants (Figs. 6-7) involving translocation of rDNA and the associated telomeres from one NOR to the other have also occurred in *fas2-4* mutants. We did not carry out mapping studies in these mutants. Note that *FAS1* and *FAS2* genes encode subunits of heterotrimeric CAF-1 (See Introduction).

### rDNA instability patterns influence rRNA variant expression patterns in *caf-1* mutants

To determine whether the altered rRNA gene variant profile at the gDNA level determine rRNA gene variant profile at RNA level, we performed RT-PCR analysis in different *fas* mutants, including *fas1-4(ΔNOR2-TEL2N), fas1-4(ΔNOR4-TEL4N), fas1-4(no NT loss*), *fas2-4(ΔNOR2-TEL2N), fas2-4(ΔNOR4-TEL4N), and fas2-4(no NT loss*. We found that the varying rRNA variant profile at the RNA level largely reflects the varying rRNA gene content at the gDNA level in *fas1-4* mutants (Fig. 8). For example, plants show relatively reduced VAR2 content at the DNA level as well as at the RNA level (Fig. 8a, first and third panel of gel images; compare relative band intensities of VAR1 to VAR2 genes among lane1 and the other lanes). Similarly, in *fas2-4* mutants, the rRNA gene variant profile at the RNA level mirrored their profile at the DNA level (Fig. 8b, compare relative band intensities of rRNA variants in the first and the third panel of gel images among lane 1 and the other lanes).

**Fig. 8.**
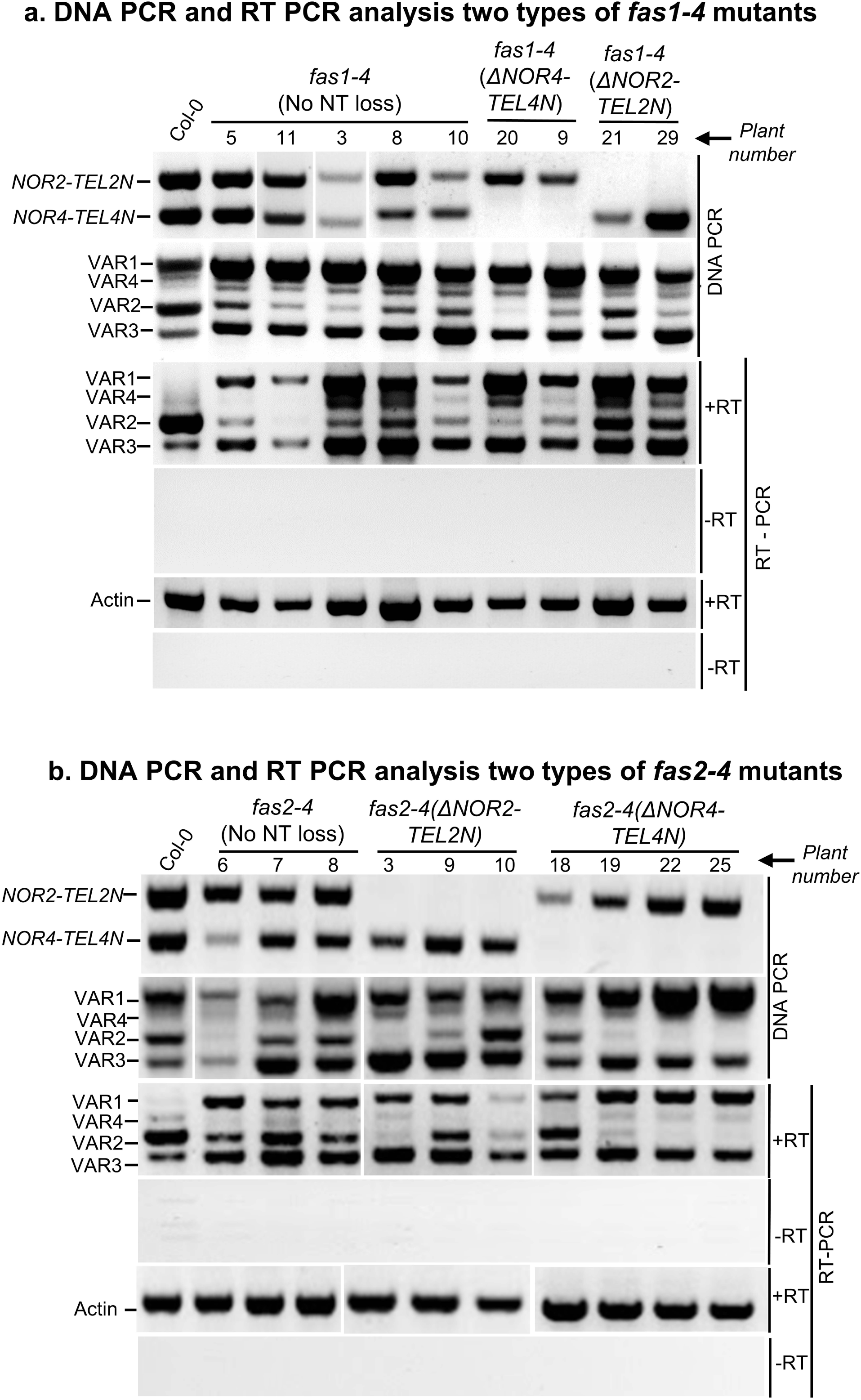
rRNA variant expression pattern in fas mutants reflects the rRNA variant pattern at the DNA level. Panels of gel images show PCR analysis of gDNA for NOR-telomere junctions and rRNA variants and, RT-PCR analysis of rRNA variants and actin (as a control) in WT Col-0 and **(a)** *fas1-4* mutants or **(b)** fas2-4 mutants.

### *cmt2* mutants did not display observable rDNA genomic instability despite the derepressed *NOR2* gene silencing

Next, we studied the effects of CMT2 loss on rDNA genomic stability. CMT2 is a plant-specific chromomethylase, which is responsible for RdDM-independent CHH methylation in Arabidopsis (17). Among the available mutant alleles, we chose *cmt2-3* (hereafter referred to as *cmt2*), a mutant carrying a T-DNA insertion (Fig. S21a), because it displayed the highest loss of CHH methylation among different *cmt2* mutant alleles (17). In a recent study, *cmt2-3* mutants displayed significant disruption of *NOR2* gene silencing (13). The study also highlighted the highest cytosine frequency in the CHH context in 45S rRNA gene regulatory elements and gene bodies. We screened ∼100 individual *cmt2* mutants for genomic instability of rDNA and its associated telomeres, as described above. Despite their significant effect on the release of *NOR2* gene silencing (Fig. 9), we did not find any observable genomic instability of rDNA or its associated telomeres in them (Fig. S21b). We genotyped all these individuals to ensure that they are *cmt2-3* mutants (Fig. S22). These findings rule out that the significant expression of NOR2-specific VAR1 genes found in the *cmt2* mutants is not influenced by any rDNA genomic instability-mediated translocation of rRNA genes from *NOR2* to *NOR4*, akin to the occurrence of such phenomena in *ddm1-2* (Figs. 2 and 3), *fas1-4* (Figs. 5-7), and *atxr5 atxr6* mutants (9).

**Fig 9:**
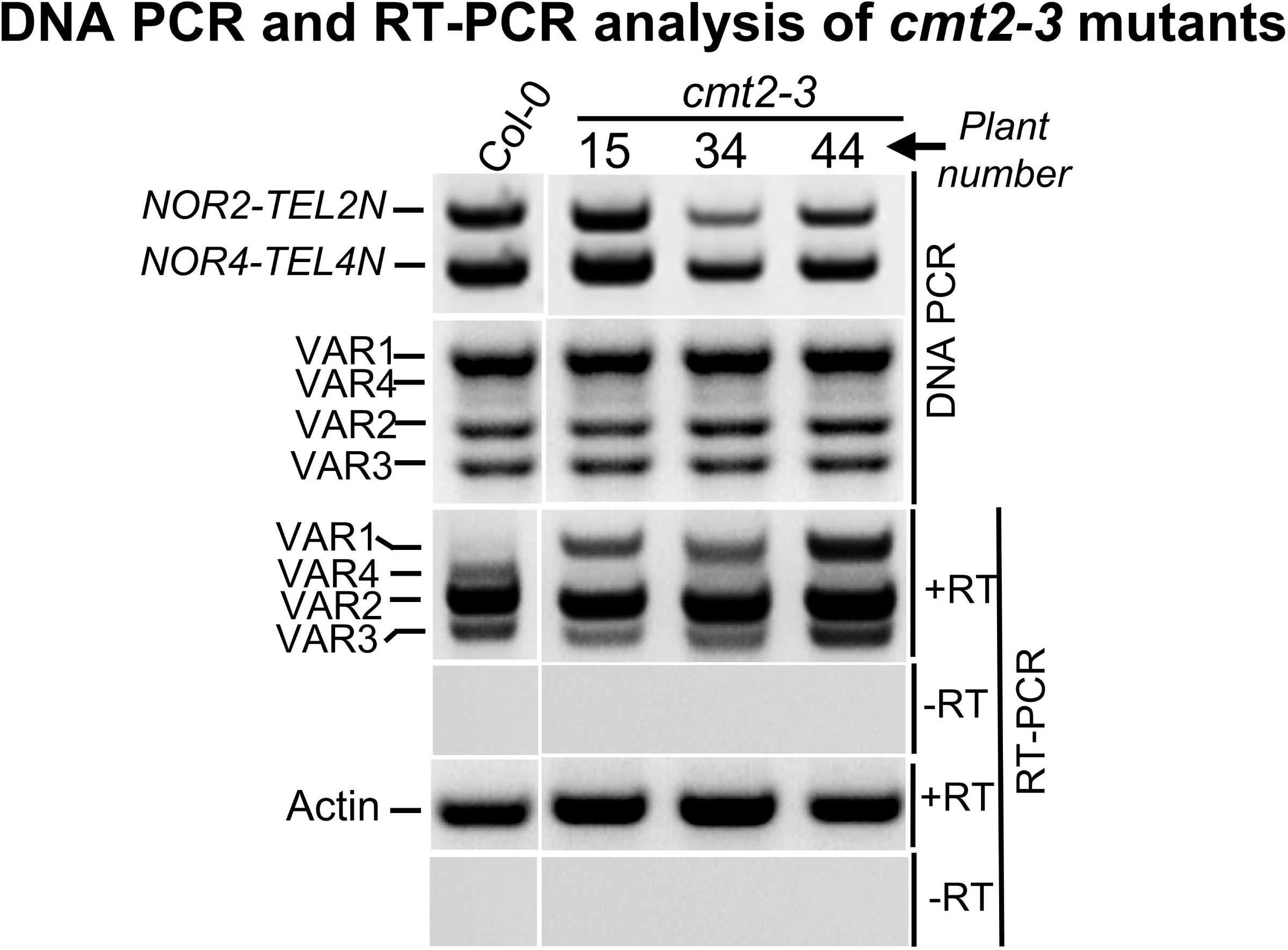
The *cmt2-3* mutants show release of rRNA gene silencing without any observable rDNA instability. The gel images show PCR analysis of gDNA for NOR-telomere junctions and rRNA variants and, RT-PCR analysis of rRNA variants and actin (as a control) in WT Col-0 and *cmt2-3* mutants.

## Discussion

Analysis of *ddm1*, *caf-1*, and *cmt2* mutants revealed the existence of multiple molecular NOR mutant phenotypes, which we named them *Mutant Phenotypes* (MP) I through IV (Fig. 10). Collectively, the data revealed that in multiple epigenetic mutants, the phenomena of rRNA gene translocation from one chromosome (NOR) to the other is occurring, which influences rRNA variant expression pattern. The *NOR2* rRNA variants are developmentally silenced in WT plants, and their expression in mutants has been interpreted as the release of *NOR2* gene silencing. Therefore, it is important to delineate the effect of an epigenetic mutation on rDNA instability and its effect in altering rRNA variant expression pattern from its effect on the actual release of rRNA gene silencing. We demonstrated such a delineation of confounding effects of multiple epigenetic mutations using genetic and molecular approaches.

**Fig. 10.**
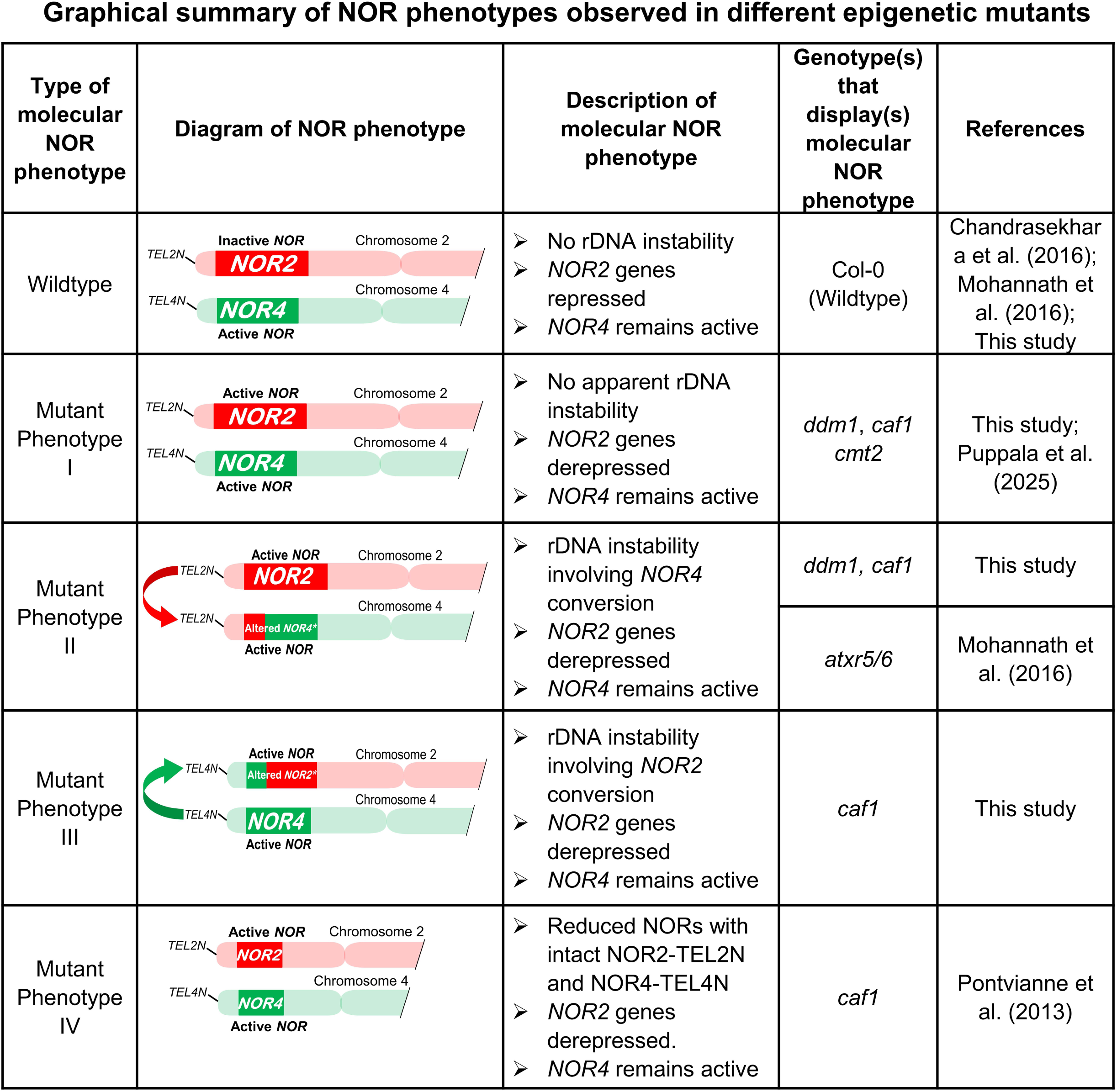
Graphical summary of different mutant NOR phenotypes observed in different epigenetic mutants in this study and in previous studies.

Based on our findings, we conclude that functions of DDM1, CAF-1, and CMT2 are required for rRNA gene silencing, and those of DDM1 and CAF-1 for maintaining rDNA genomic stability. DDM1 is required for DNA methylation in all three cytosine sequence contexts (CG, CHG, and CHH) and heterochromatinization (17, 25). Mechanistically, DDM1 has been demonstrated to deposit histone 2A variant W (H2A.W) onto mobile transposable elements (TEs), resulting in their heterochromatinization (24). Moreover, DDM1 also requires CAF-1 for H3.1 deposition for heterochromatinization (23). These studies reveal why deficiency of CAF-1, DDM1 or CMT2 results in a release of rRNA gene silencing.

Compared to both *caf-1* (*fas1-4* and *fas2-4*) and *ddm1-2* mutants, we could not detect any observable rDNA genomic instability in the *ddm1-1* and *cmt2-3* mutants, despite both mutations resulting in a significant release of rRNA gene silencing. Nonetheless, we do not rule out the occurrence of rDNA instability in these mutants, which display loss of DNA methylation (17, 38). The reasoning is that, given the stochastic nature of the rDNA genomic instabilities observed so far, screening a further large number of plants or multiple generations may reveal some rDNA instability in these mutants as well.

The *ddm1-2* mutants carry a truncated DDM1 protein, deficient for two functional domains; a conserved H2A.W-binding domain and a HELICc domain (Fig. S12c) (23, 24), while *ddm1-1* mutants carry a single amino acid substitution in the HELICc domain (Fig. S12b), which reduces DDM1 functionality. Therefore, it is not surprising to observe a more severe mutant phenotype in *ddm1-2* compared to *ddm1-1* mutants. However, the loss of rRNA silencing in DNA methylation-defective *ddm1-1* mutants (38), further highlights the importance of the HELICc domain in DDM1 protein function in cytosine methylation and gene silencing. In *ddm1-1* mutants, cysteine to tyrosine substitution at the 615^th^ position is predicted to specifically disrupt the disulfide bond (S-S) between two cysteine residues at positions 615 and 634 (23). DNA methylation plays a major role in heterochromatinization-mediated silencing, but a previous study demonstrated that gene silencing could also occur independently of DNA methylation (39).

Likewise, the fact that CAF-1 deposits histone H3.1 required for heterochromatin formation (27, 40, 41), explains the severity of the *caf-1* mutant phenotype encompassing severe loss of rDNA and significant release of rRNA gene silencing (Figs. 5-7) (33, 34) when compared to that of *atxr5 atxr6* mutants, which displayed a *NOR4* conversion and a modest release of rRNA gene silencing (9). Note that ATXR5/6 monomethylates lysine 27 of histone H3.1 (H3.1K27me1), which CAF-1 deposits for heterochromatin formation, and H3.1K27me1 is one of the several heterochromatin marks that histone H3.1 carries (35, 42, 43). Thus, the mutant phenotype of *caf-1* is predicted to be more severe than the *atxr5 atxr6* mutants.

DNA cytosine methylation has been demonstrated to inhibit homologous recombination in multiple eukaryotes and inferred to play a role in preserving genome integrity (44–46). In Arabidopsis, the loss of DDM1 affects the recombination landscape without affecting the total number of crossovers (47, 48). Furthermore, the loss of DDM1 inhibits the homology-directed repair of DNA double-strand breaks (DSBs) (49). Similarly, homology-dependent repair has been implicated in rDNA loss in *caf-1* mutants, which accumulate more DSBs than the wild-type plants (50). However, NOR conversion events involving translocation of rDNA from one NOR to the other, found in the *ddm1* and *caf-1* mutants, may not be explained by the dysregulated homologous recombination events. More than one mechanism could account for the translocation of rDNA from one NOR to the other. One possibility is that a nonreciprocal translocation of rRNA genes from one NOR to the other resulted from a break-induced replication (51–54), initiated by a collapsed DNA replication fork or double-strand break within one of the NORs. Homologous repair initiated by one broken strand of the NOR invading the essentially identical sequences of the other NOR, instead of using the same NOR sequences of the presumably unbroken homologous (sister) chromosome, followed by branch migration of the resulting D-loop till the end of the chromosome, could potentially account for replacement of the lost NOR and the associated telomere sequences by the corresponding sequences from the other NOR (Fig. S23). Such a possibility has also been speculated for *NOR4* conversion observed in *atxr5 atxr6* mutants (9). The other possibility is a recombination event involving a single non-homologous crossover between chromosomes 2 and 4 occurring within the NORs, followed by independent segregation of crossover products.

Whatever may be the mechanism of NOR conversion, this study highlights the salient roles of chromatin remodelling, including DNA methylation and various histone modifications, in maintaining rDNA genomic stability and rRNA gene silencing. The drastic genomic instabilities observed in these mutants seem to be limited to NOR regions, but a thorough analysis of the whole genome sequences is required to determine the extent of stochastic genomic instability in the remainder of the genome, if any.

## Acknowledgements

We thank Prof. Craig S. Pikaard Howard Hughes Medical Institute, Indiana University, USA, for generously sharing seeds of several Arabidopsis mutants and ecotypes. We thank members of the Department of Biological Sciences and other divisions, BITS-Pilani Hyderabad campus, for their help.

## Author contribution statement

M.K.R., A.T.S., R.T., and G.M. designed research; M.K.R., A.T.S., R.T., A.B., B.A., S.G., G.P.S., and S.R., performed research; M.K.R., A.T.S., R.T., and G.M. analyzed data; M.K.R., A.T.S., R.T., and G.M. wrote the paper.

## References

1. J. Baßler, E. Hurt, Eukaryotic Ribosome Assembly. Annu Rev Biochem 88, 281–306 (2019).

2. J. Sáez-Vásquez, M. Delseny, Ribosome Biogenesis in Plants: From Functional 45S Ribosomal DNA Organization to Ribosome Assembly Factors. The Plant cell 31, 1945–1967 (2019).

3. T. W. Turowski, D. Tollervey, Cotranscriptional events in eukaryotic ribosome synthesis. Wiley interdisciplinary reviews. RNA 6, 129–139 (2015).

4. O. V. Viktorovskaya, D. A. Schneider, Functional divergence of eukaryotic RNA polymerases: unique properties of RNA polymerase I suit its cellular role. Gene 556, 19–26 (2015).

5. G. P. Copenhaver, C. S. Pikaard, Two-dimensional RFLP analyses reveal megabase-sized clusters of rRNA gene variants in Arabidopsis thaliana, suggesting local spreading of variants as the mode for gene homogenization during concerted evolution. Plant J 9, 273–282 (1996).

6. G. P. Copenhaver, C. S. Pikaard, RFLP and physical mapping with an rDNA-specific endonuclease reveals that nucleolus organizer regions of Arabidopsis thaliana adjoin the telomeres on chromosomes 2 and 4. Plant J 9, 259–272 (1996).

7. D. Fultz, A. McKinlay, R. Enganti, C. S. Pikaard, Sequence and epigenetic landscapes of active and silent nucleolus organizer regions in Arabidopsis. Science advances 9, eadj4509 (2023).

8. C. Chandrasekhara, G. Mohannath, T. Blevins, F. Pontvianne, C. S. Pikaard, Chromosome-specific NOR inactivation explains selective rRNA gene silencing and dosage control in Arabidopsis. Genes Dev 30, 177–190 (2016).

9. G. Mohannath, F. Pontvianne, C. S. Pikaard, Selective nucleolus organizer inactivation in Arabidopsis is a chromosome position-effect phenomenon. Proceedings of the National Academy of Sciences 113, 13426–13431 (2016).

10. F. Pontvianne et al., Nucleolin is required for DNA methylation state and the expression of rRNA gene variants in Arabidopsis thaliana. PLoS genetics 6, e1001225 (2010).

11. W. Mo et al., Single-molecule targeted accessibility and methylation sequencing of centromeres, telomeres and rDNAs in Arabidopsis. Nat Plants 9, 1439–1450 (2023).

12. G. Mohannath, C. S. Pikaard, Analysis of rRNA Gene Methylation in Arabidopsis thaliana by CHEF-Conventional 2D Gel Electrophoresis. Methods in molecular biology (Clifton, N.J.) 1455, 183–202 (2016).

13. N. V. Puppala, R. Tammineni, G. P. Saradadevi, G. Mohannath, Combination of CG methylation and Chromomethylase 2-mediated RdDM-independent CHH methylation is required for chromosome-specific rRNA gene silencing. bioRxiv 10.1101/2023.02.03.526984, 2023.2002.2003.526984 (2026).

14. F. Pontvianne et al., Histone methyltransferases regulating rRNA gene dose and dosage control in Arabidopsis. Genes Dev 26, 945–957 (2012).

15. S. J. Cokus et al., Shotgun bisulphite sequencing of the Arabidopsis genome reveals DNA methylation patterning. Nature 452, 215–219 (2008).

16. R. Lister et al., Highly integrated single-base resolution maps of the epigenome in Arabidopsis. Cell 133, 523–536 (2008).

17. A. Zemach et al., The Arabidopsis nucleosome remodeler DDM1 allows DNA methyltransferases to access H1-containing heterochromatin. Cell 153, 193–205 (2013).

18. J. A. Law, S. E. Jacobsen, Establishing, maintaining and modifying DNA methylation patterns in plants and animals. Nature reviews. Genetics 11, 204–220 (2010).

19. B. Rymen, L. Ferrafiat, T. Blevins, Non-coding RNA polymerases that silence transposable elements and reprogram gene expression in plants. Transcription 11, 172–191 (2020).

20. E. Sasaki, T. Kawakatsu, J. R. Ecker, M. Nordborg, Common alleles of CMT2 and NRPE1 are major determinants of CHH methylation variation in Arabidopsis thaliana. PLoS genetics 15, e1008492 (2019).

21. A. J. Bewick et al., The evolution of CHROMOMETHYLASES and gene body DNA methylation in plants. Genome biology 18, 65 (2017).

22. J. Brzeski, A. Jerzmanowski, Deficient in DNA methylation 1 (DDM1) defines a novel family of chromatin-remodeling factors. The Journal of biological chemistry 278, 823–828 (2003).

23. S. C. Lee et al., Chromatin remodeling of histone H3 variants by DDM1 underlies epigenetic inheritance of DNA methylation. Cell 186, 4100–4116 e4115 (2023).

24. A. Osakabe et al., The chromatin remodeler DDM1 prevents transposon mobility through deposition of histone variant H2A. W. Nature cell biology 23, 391–400 (2021).

25. J. A. Jeddeloh, T. L. Stokes, E. J. Richards, Maintenance of genomic methylation requires a SWI2/SNF2-like protein. Nature genetics 22, 94–97 (1999).

26. A. Vongs, T. Kakutani, R. A. Martienssen, E. J. Richards, Arabidopsis thaliana DNA methylation mutants. Science 260, 1926–1928 (1993).

27. E. Ramirez-Parra, C. Gutierrez, The many faces of chromatin assembly factor 1. Trends in plant science 12, 570–576 (2007).

28. S. Smith, B. Stillman, Purification and characterization of CAF-I, a human cell factor required for chromatin assembly during DNA replication in vitro. Cell 58, 15–25 (1989).

29. H. Tagami, D. Ray-Gallet, G. Almouzni, Y. Nakatani, Histone H3.1 and H3.3 complexes mediate nucleosome assembly pathways dependent or independent of DNA synthesis. Cell 116, 51–61 (2004).

30. L. Hennig, P. Taranto, M. Walser, N. Schönrock, W. Gruissem, Arabidopsis MSI1 is required for epigenetic maintenance of reproductive development. Development 130, 2555–2565 (2003).

31. H. Kaya et al., FASCIATA genes for chromatin assembly factor-1 in arabidopsis maintain the cellular organization of apical meristems. Cell 104, 131–142 (2001).

32. H. M. O. Leyser, I. J. Furner, Characterisation of three shoot apical meristem mutants of Arabidopsis thaliana. Development 116, 397–403 (1992).

33. I. Mozgova, P. Mokros, J. Fajkus, Dysfunction of chromatin assembly factor 1 induces shortening of telomeres and loss of 45S rDNA in Arabidopsis thaliana. The Plant cell 22, 2768–2780 (2010).

34. F. Pontvianne et al., Subnuclear partitioning of rRNA genes between the nucleolus and nucleoplasm reflects alternative epiallelic states. Genes & development 27, 1545–1550 (2013).

35. Y. Jacob et al., Selective methylation of histone H3 variant H3.1 regulates heterochromatin replication. Science 343, 1249–1253 (2014).

36. R. Yadegari et al., Mutations in the FIE and MEA genes that encode interacting polycomb proteins cause parent-of-origin effects on seed development by distinct mechanisms. The Plant cell 12, 2367–2381 (2000).

37. G. P. Saradadevi et al., Structural variation among assembled genomes facilitates development of rapid and low-cost NOR-linked markers and NOR-telomere junction mapping in Arabidopsis. Plant cell reports 42, 1059–1069 (2023).

38. J. A. Jeddeloh, J. Bender, E. J. Richards, The DNA methylation locusDDM1 is required for maintenance of gene silencing in Arabidopsis. Genes & development 12, 1714–1725 (1998).

39. E. Hristova, K. Fal, L. Klemme, D. Windels, E. Bucher, HISTONE DEACETYLASE6 Controls Gene Expression Patterning and DNA Methylation-Independent Euchromatic Silencing. Plant physiology 168, 1298–1308 (2015).

40. A. Kirik, A. Pecinka, E. Wendeler, B. Reiss, The chromatin assembly factor subunit FASCIATA1 is involved in homologous recombination in plants. The Plant cell 18, 2431–2442 (2006).

41. N. Schönrock, V. Exner, A. Probst, W. Gruissem, L. Hennig, Functional genomic analysis of CAF-1 mutants in Arabidopsis thaliana. The Journal of biological chemistry 281, 9560–9568 (2006).

42. Y. Jacob et al., ATXR5 and ATXR6 are H3K27 monomethyltransferases required for chromatin structure and gene silencing. Nat Struct Mol Biol 16, 763–768 (2009).

43. T. Zhao, Z. Zhan, D. Jiang, Histone modifications and their regulatory roles in plant development and environmental memory. J Genet Genomics 46, 467–476 (2019).

44. L. Maloisel, J. L. Rossignol, Suppression of crossing-over by DNA methylation in Ascobolus. Genes Dev 12, 1381–1389 (1998).

45. N. E. Yelina et al., DNA methylation epigenetically silences crossover hot spots and controls chromosomal domains of meiotic recombination in Arabidopsis. Genes Dev 29, 2183–2202 (2015).

46. N. Zamudio et al., DNA methylation restrains transposons from adopting a chromatin signature permissive for meiotic recombination. Genes Dev 29, 1256–1270 (2015).

47. M. Colome-Tatche et al., Features of the Arabidopsis recombination landscape resulting from the combined loss of sequence variation and DNA methylation. Proc Natl Acad Sci U S A 109, 16240–16245 (2012).

48. C. Melamed-Bessudo, A. A. Levy, Deficiency in DNA methylation increases meiotic crossover rates in euchromatic but not in heterochromatic regions in Arabidopsis. Proc Natl Acad Sci U S A 109, E981–988 (2012).

49. S. H. Choi et al., Mutation in DDM1 inhibits the homology directed repair of double strand breaks. PloS one 14, e0211878 (2019).

50. V. Muchová et al., Homology-dependent repair is involved in 45 S rDNA loss in plant CAF-1 mutants. The Plant Journal 81, 198–209 (2015).

51. R. P. Anand, S. T. Lovett, J. E. Haber, Break-induced DNA replication. Cold Spring Harbor perspectives in biology 5, a010397 (2013).

52. G. Bosco, J. E. Haber, Chromosome break-induced DNA replication leads to nonreciprocal translocations and telomere capture. Genetics 150, 1037–1047 (1998).

53. P. Hastings, Mechanisms of ectopic gene conversion. Genes 1, 427–439 (2010).

54. N. Saini et al., Migrating bubble during break-induced replication drives conservative DNA synthesis. Nature 502, 389–392 (2013).

