## Supplementary Figures for "Genomic rDNA instabilities in Arabidopsis epigenetic mutants alter location-based rRNA gene expression patterns"

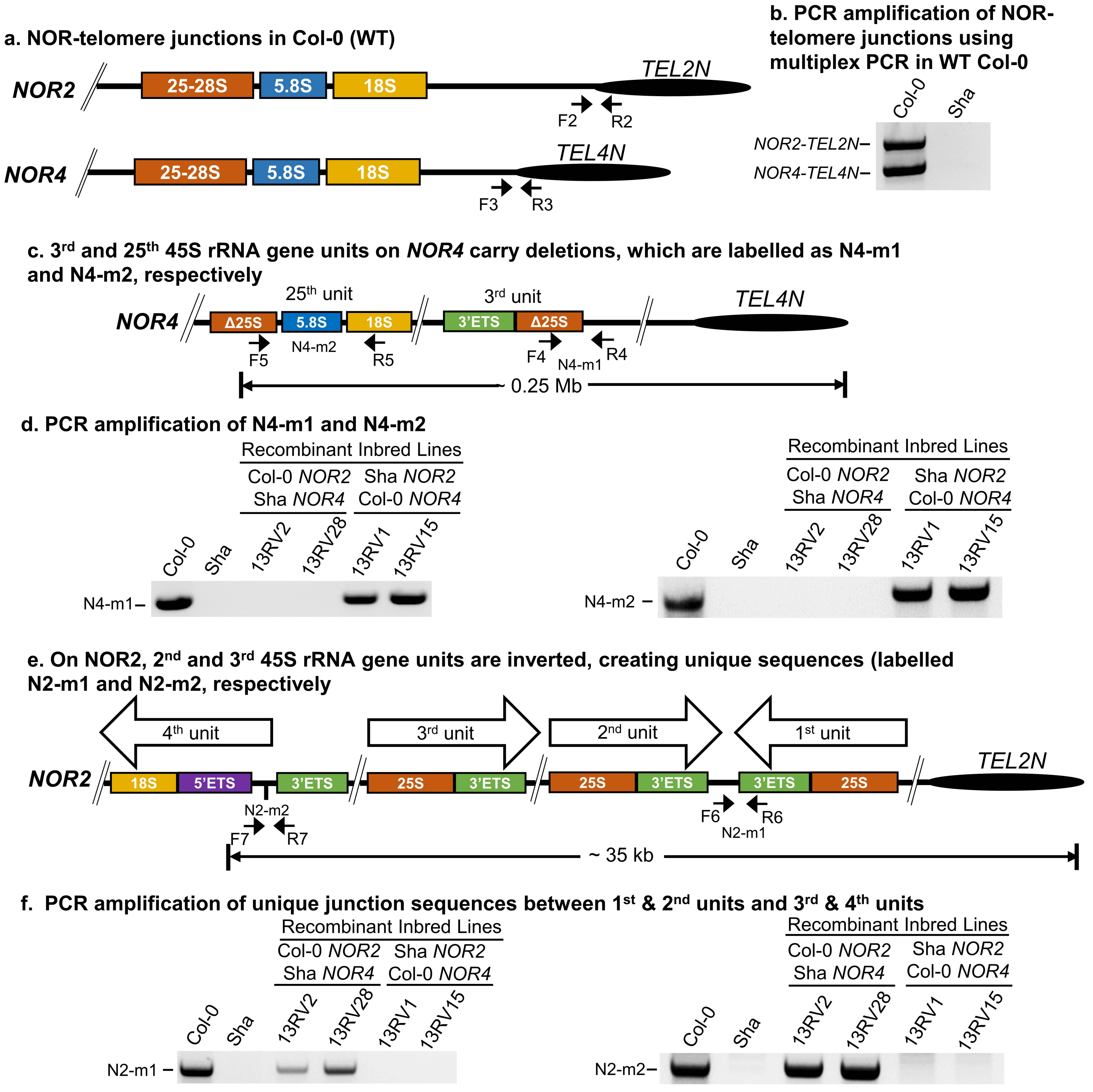


**Figure S1:** Several PCR-based markers were designed to amplify unique regions of *NOR2* and *NOR4* in Col-0. **a** Diagram depicts the *NOR2-TEL2N* and the *NOR4-TEL4N* junction sequences of WT Col-0. Also shown are the relative positions of primer pairs used to amplify the NOR-telomere junctions. **b** The gel image shows PCR amplification of NOR-Telomere junctions in Col-0, and Sha is used as a negative control. **c** Diagram shows the 3rd and 25th 45S rRNA gene units in *NOR4* that carry large deletions, which are labelled as N4-m1 and N4-m2, respectively. Also shown are the relative positions of primer pairs used to amplify N4-m1 and N4-m2 markers. **d** The gel images show PCR amplification of N4-m1 and N4-m2 in WT Col-0, Sha, and RILs of Col-0 x Sha cross, which carry Col-0 *NOR2* and Sha *NOR4* or Col-0 *NOR4* and Sha *NOR2*. **e** Diagram shows the first four 45S rRNA gene units, of which, 2nd and 3rd unit are inverted. The unique junction sequences between 1st and 2nd unit, and 3rd and 4th are labelled as N2-m1 and N2-m2, respectively. Also shown are the relative positions of primer pairs used to amplify N2-m1 and N2-m2 markers. **f** The gel images show PCR amplification of N2-m1 and N2-m2 in WT Col-0, Sha, and RILs of Col-0 x Sha cross, which carry Col-0 *NOR2* and Sha *NOR4* or Col-0 *NOR4* and Sha *NOR2*.

**a. *ddm1-2* mutant has a G→ A substitution at splice donor site of intron 11**

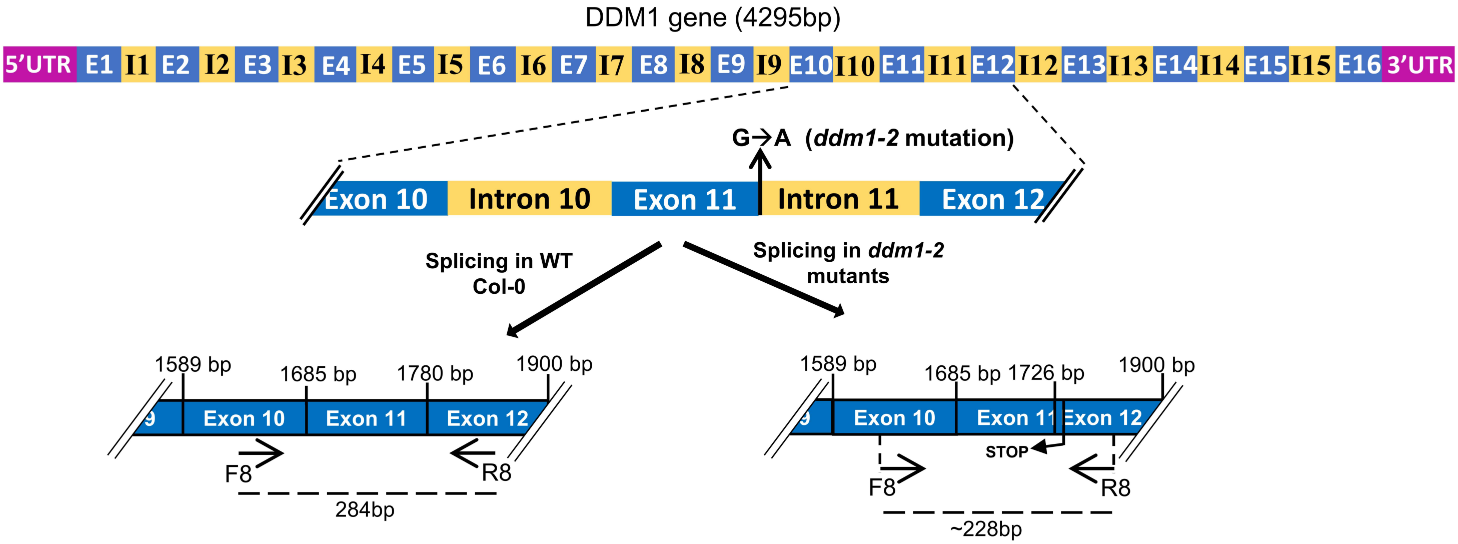


**b. Defective splicing in *ddm1-2* mutants causes a deletion of 54bp in exon 11 and 2 bp in exon 12, leading to a frameshift mutation and premature translation termination**

Col-0 (Wildtype)

**Forward primer (F8)**

**GCUGGAAGGGAAAGCUUAACAACC**UGGUCAUU
CAACUUCGAAAGAACUGCAACCAUCCUGACCU
UCUCCAGGGGCAAAUAGAUGGUUCAUAUCUCU
ACCCUCCUGUUGAAGAGAUUGUUGGACAGUGU
GGUAAAUUCCGCUUAUUGGAGAGAUUACUUGU
UCGGUUAUUUGCCAAUAAUCACAAAGUCCUUA
UCUUCUCCCAAUGGACGAAACUUUUGGACAUU
AUGGAUUACUACUUCAGUGAGAAGGGGUUUGA
**GGUUUGCAGAAUCGAUGGCAGUGU**

**Reverse primer (R8)**

*ddm1-2*

**Forward primer (F8)**

**GCUGGAAGGGAAAGCUUAACAACC**UGGUCAUU
CAACUUCGAAAGAACUGCAACCAUCCUGACCU
UCUCCAGGGGCAAAUAGAUGGUUCAUAUCUCU
ACCCUCCUGUUGAAGAGAUUGUUGGACAGUGU
GGU-----------------------------
---------------------------CCUUA
UCUUCUCCCAAUGGACGAAACUUUUGGACAUU
AUGGAUUACUACUUCAGUGAGAAGGGGUUUGA
**GGUUUGCAGAAUCGAUGGCAGUGU**

**Reverse primer (R8)**

**c. PCR and RT-PCR analysis of the mis-spliced region**

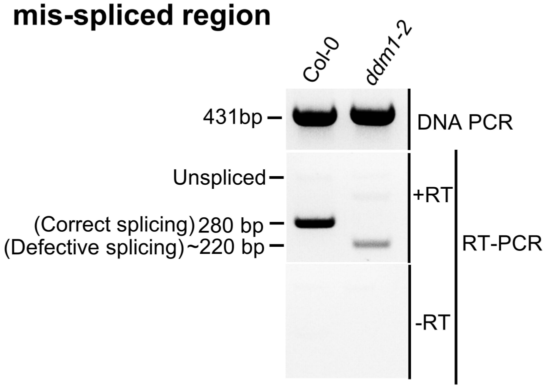


**Figure S2**: G to A substitution in *ddm1-2* mutants causes defective splicing leading introduction of a premature stop codon. **(a)** Cartoon shows location of G to A substitution in *DDM1* gene and its effect on splicing. **(b)** A diagram depicts the deleted mRNA sequences in *ddm1-2* mutants compared to WT-Col-0. **(c)** A gel image shows PCR and RT-PCR analysis of gDNA and cDNA, respectively, for the defectively spliced region in WT-Col-0 and *ddm1-2* mutants.

#### a. Illustration of *DDM1* region used for dCAPS assay to identify *ddm1-2* mutants

#### b. Genotyping of *ddm1-2* mutants by dCAPS assay

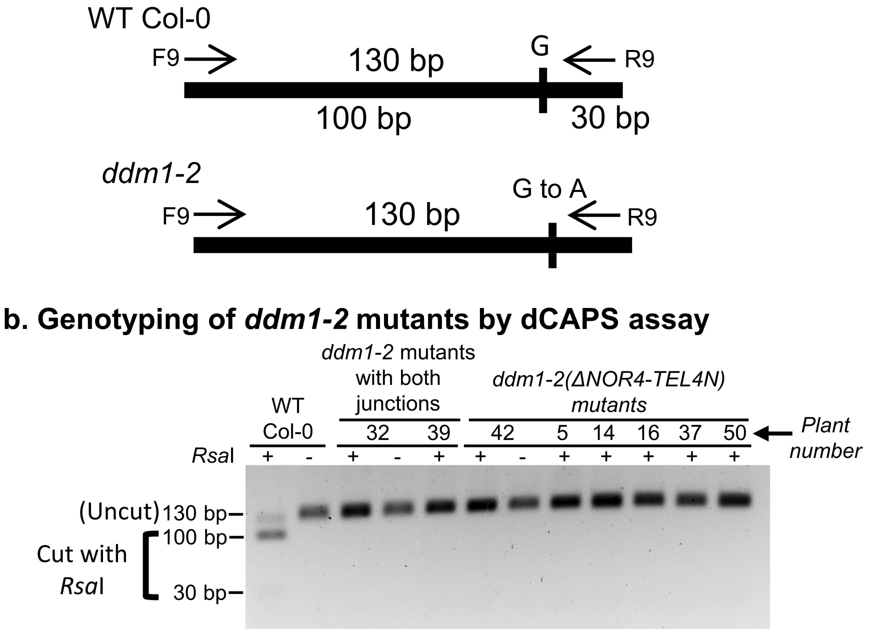


**Figure S3** Genotyping of *ddm1-2* mutants. **a** A line diagram shows the site the G to A substitution in *ddm1-2* mutants, which was exploited to design a dCAPS assay, which introduces a *Rsa*I site in WT Col-0 amplicons but not in those of *ddm1-2* mutants (Yadegari et al., 2000). **b** A gel image shows dCAPS assay of WT Col-0 and *ddm1-2* mutants.

**a Diagrammatic representation of NOR4-adjacent markers**

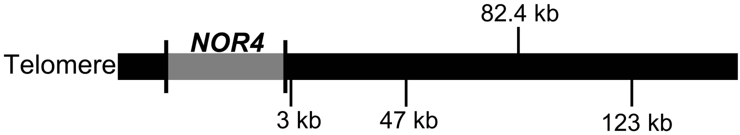


**b PCR amplification of NOR4-adjacent markers**

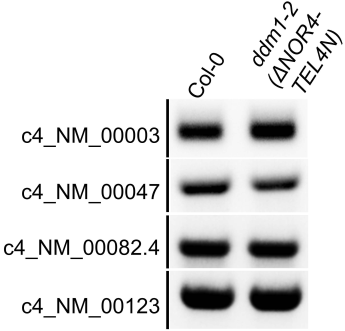


**Figure S4**: NOR4-adjoining regions are intact in *ddm1-2* mutants. **a** A cartoon depicts NOR4-adjacent markers. **b** The gel image shows PCR amplification NOR4-adjacent markers in *ddm1-2* mutants and WT Col-0.

#### PCR analysis of NOR-telomere junctions, *NOR4* markers (N4-m1 and N4-m2), and rRNA variants

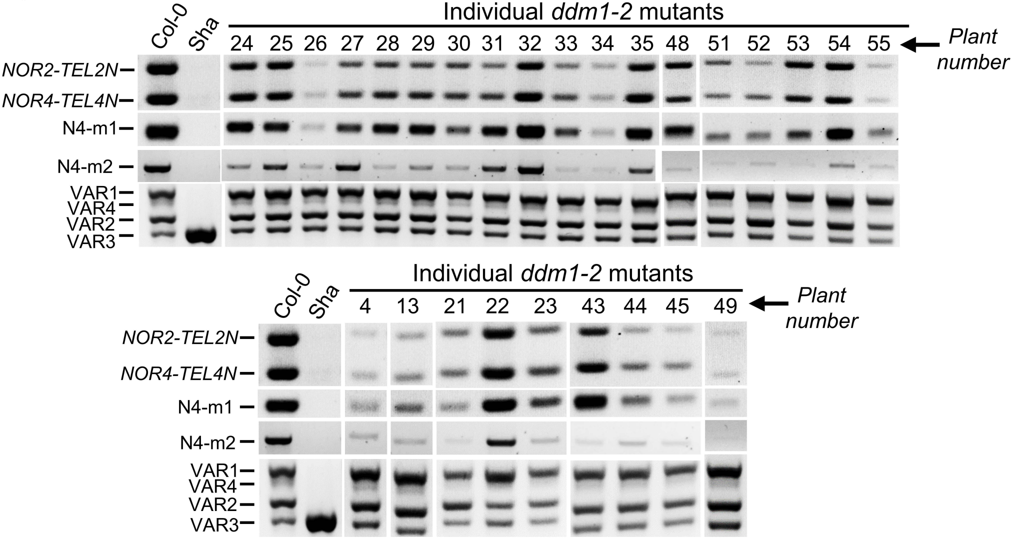


**Figure S5** Characterization of genomic instability pattern of 45S rRNA genes and the associated telomeres in the remainder of *ddm1-2* mutants The gel images show PCR analysis of NOR-telomere junctions, *NOR4* markers (N4-m1 and N4-m2), and rRNA gene subtypes, in WT-Col-0, Sha, and the remainder of individual *ddm1-2* mutant plants from a set of 52. The remaining are shown in Fig. 2.

#### PCR analysis of NOR-telomere junctions in *ddm1-2* mutants in $G_{n+1}$ generation

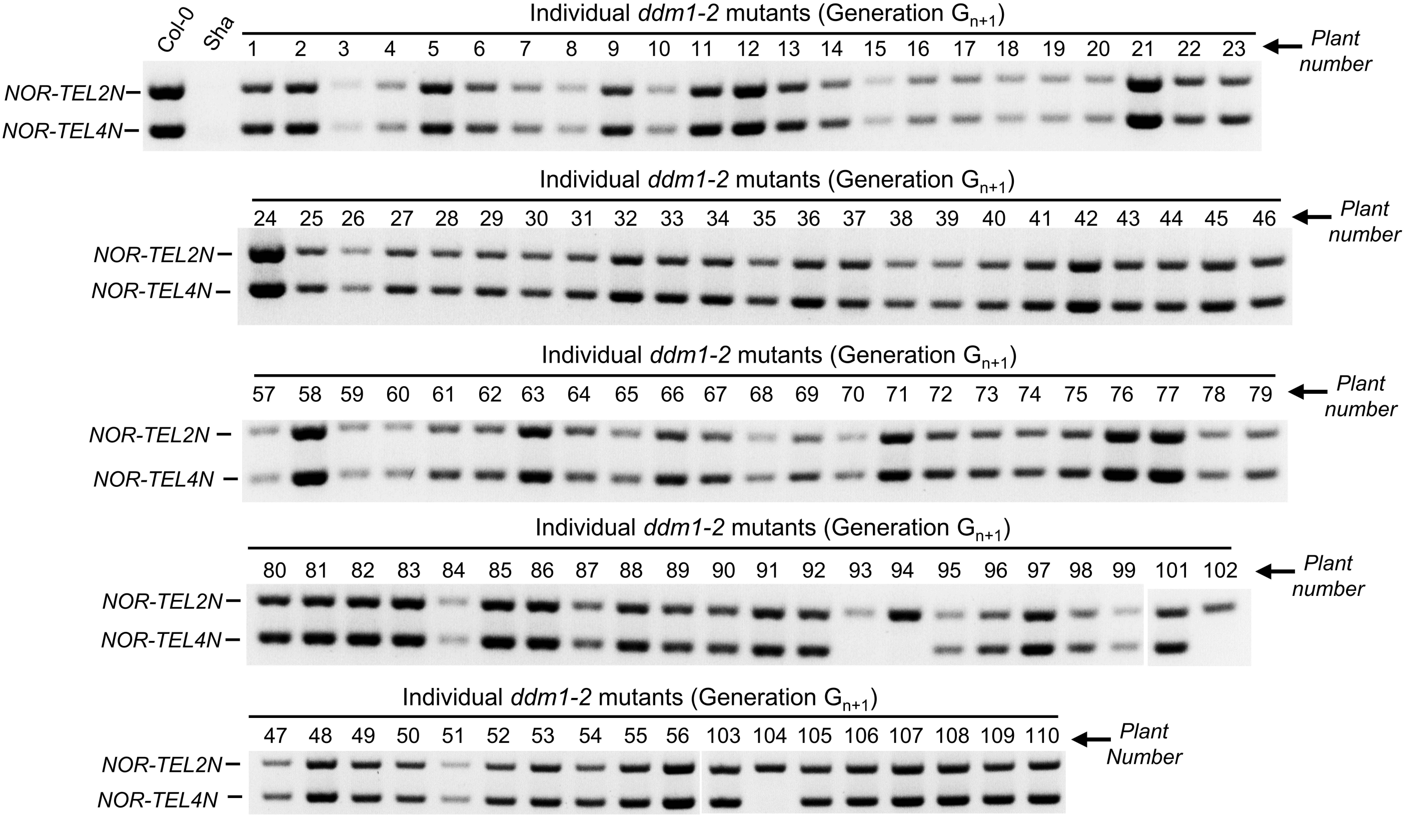


**Figure S6** The genomic instabilities are mainly associated with *NOR4* and the associated telomere. The gel images show PCR amplification of NOR-telomere junctions in WT Col-0, Sha, and *ddm1-2* mutants from the $G_{n+1}$ generation.

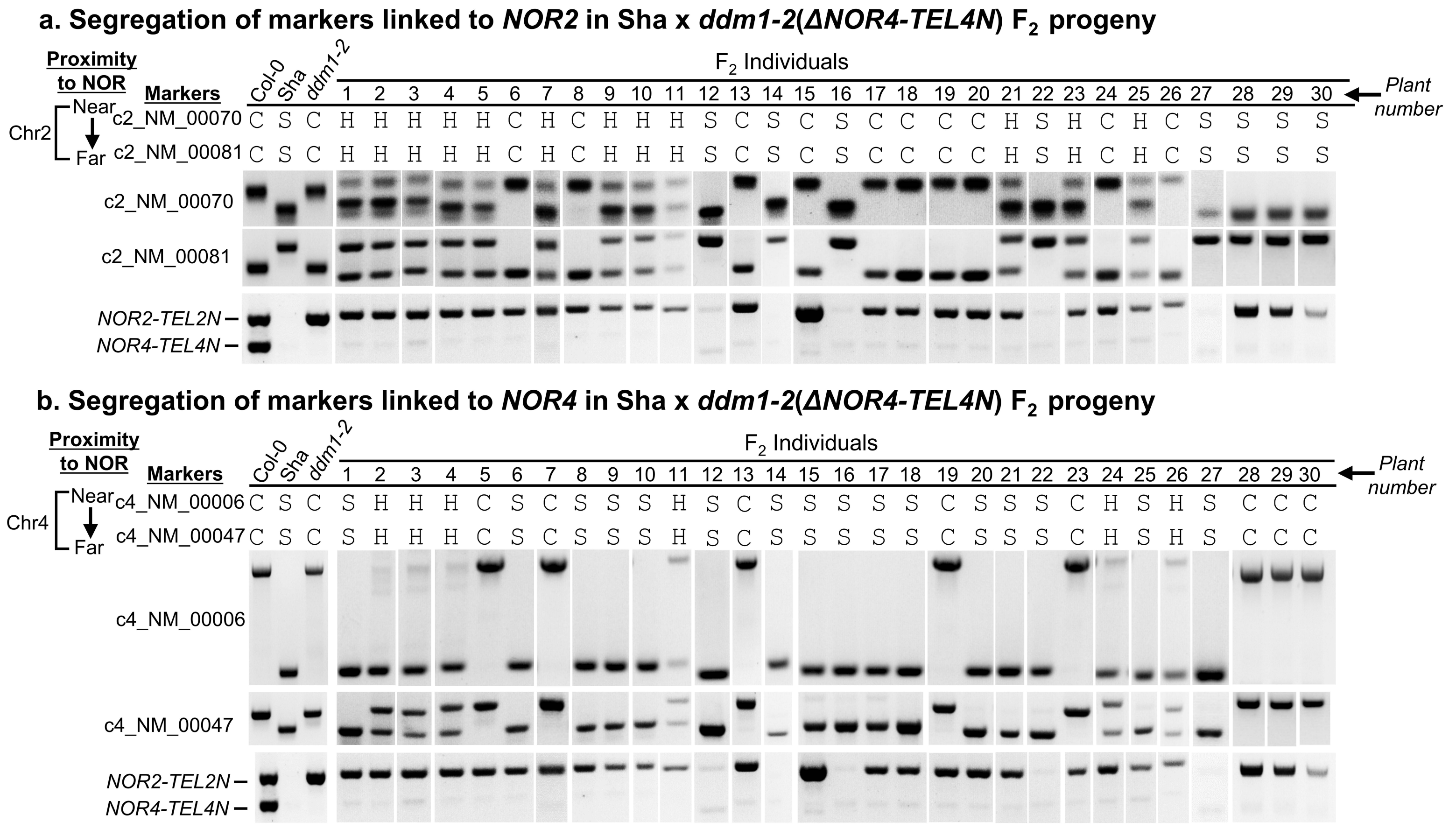


**Figure S7**: Genotyping of Sha x *ddm1-2(ΔNOR4-TEL4N)* $F_2$ progeny. The panels show gel images of length polymorphism-based PCR markers that distinguish Col-0 and Sha alleles in the regions adjacent to *NOR2* and *NOR4* **(a)** Segregation of markers linked to *NOR2.* **(b)** Segregation of markers linked to *NOR4.*
C=Col-0; S=Sha; H=heterozygous.

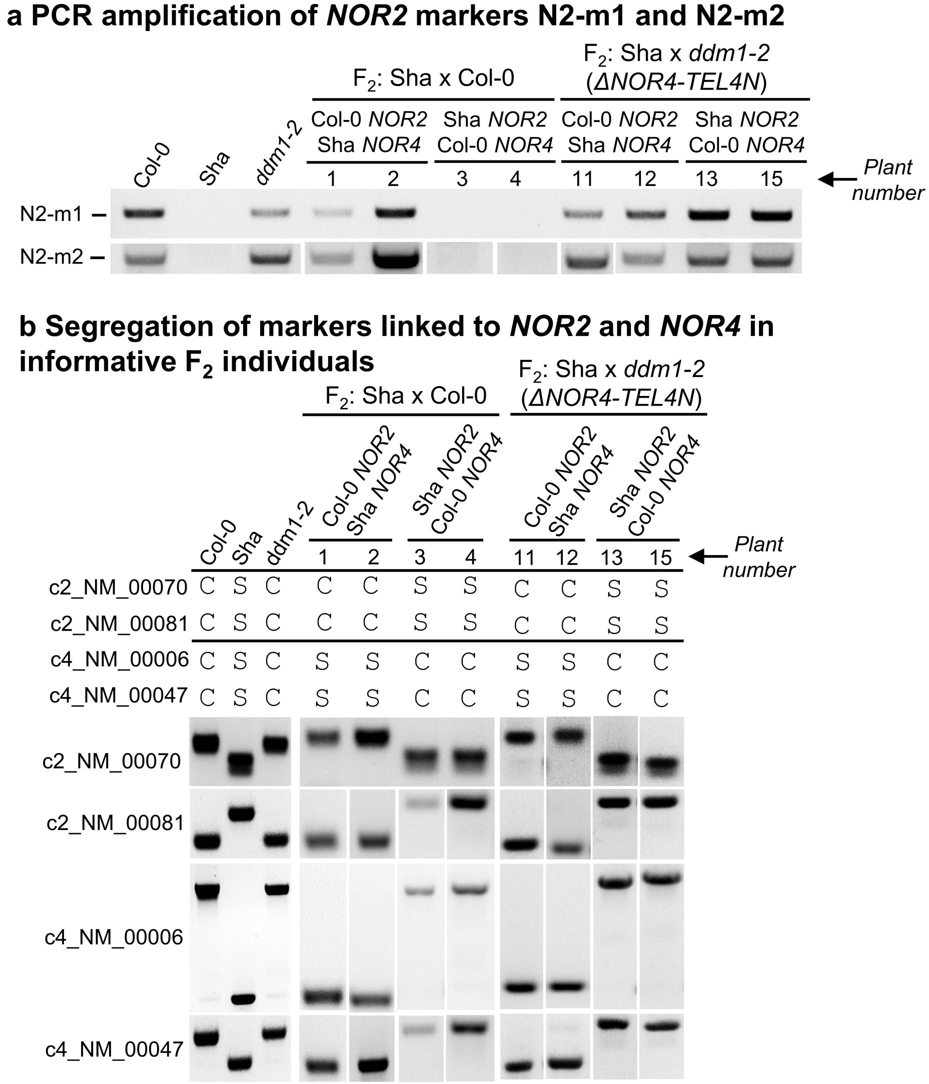


**Figure S8**: In *ddm1-2* mutants, telomere-proximal region of *NOR4,* including *NOR4-TEL4N* (at least 0.25 Mb) has been deleted and replaced by the corresponding *NOR2* sequences. Gel images show PCR amplification of **(a)** *NOR2* markers N2-m1 and N2-m2 **(b)** NOR2- and NOR4-linked markers, in informative $F_2$ individuals of Sha x Col-0 and Sha x *ddm1-2*(*ΔNOR4-TEL4N*), Col-0 and Sha controls.

**a. Chi-square test results for NOR2-linked marker c2_NM_00081**

| Genotype | Observed | Expected | χ2 | df | P(alpha) |
|---|---|---|---|---|---|
| Col-0 | 30 | 24.5 | 4.102 | 2 | Not significant |
| Heterozygous | 39 | 49 | | | |
| Sha | 29 | 24.5 | | | |
| Total number of plants genotyped: 98 | | | | | |

**b. Chi-square test results for NOR4-linked marker c4_NM_00047**

| Genotype | Observed | Expected | χ2 | df | P(alpha) |
|---|---|---|---|---|---|
| Col-0 | 26 | 24.5 | 2.326 | 2 | Not significant |
| Heterozygous | 42 | 49 | | | |
| Sha | 30 | 24.5 | | | |
| Total number of plants genotyped: 98 | | | | | |

**Figure S9**: Chi-square statistic analysis show that NOR2- and NOR4-linked markers are segregating normally in the Mendelian fashion in F2 progeny of Sha x *ddm1-2*(*ΔNOR4-TEL4N*). Tables show χ2 test results for **(a)** NOR2-linked marker c2_NM_00081 and **(b)** NOR4-linked marker c4_NM_00047, in the $F_2$ progeny of Sha x *ddm1-2*(*ΔNOR4-TEL4N*)
df=degrees of freedom; p= probability

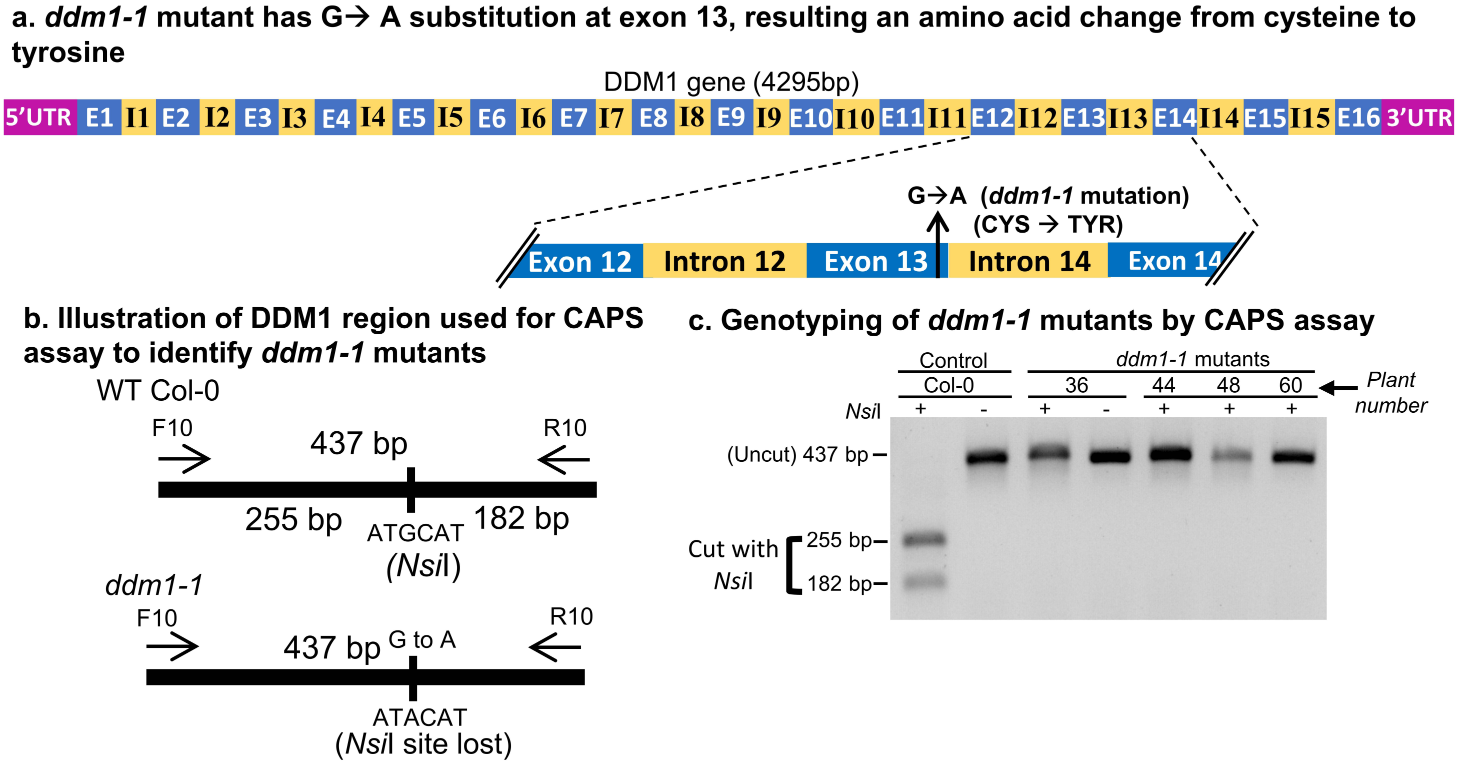


**Figure S10** Location of G to A substitution in *ddm1-1* mutants and their genotyping. **a** Cartoon shows location of G to A substitution in DDM1 gene and its effect on amino acid substitution. **b** A line diagram shows the loss of *Nsi*I site in *ddm1-1* mutants, which was used to design a CAPS assay. **c** A gel image shows CAPS assay of WT Col-0 and *ddm1-1* mutants.

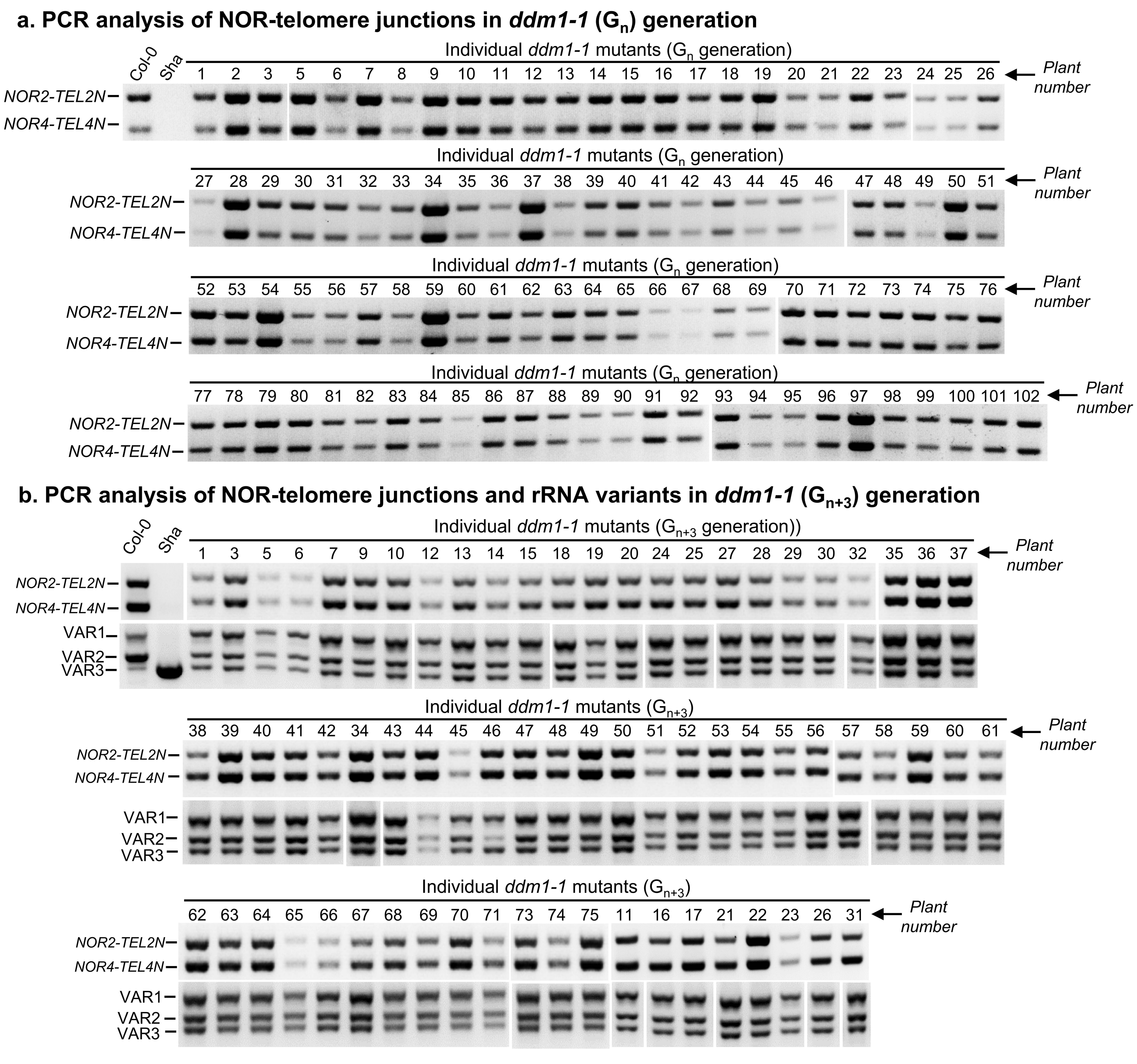


**Figure S11:** The *ddm1-1* mutants did not show any observable genomic instabilities of rDNA or the associated telomeres. **a** Panels of gel images show PCR analysis NOR-telomere junctions of *ddm1-1* mutants of ($G_n$) generation. **b** Panels of gel images show PCR analysis NOR-telomere junctions and rRNA variants of *ddm1-1* mutants of ($G_{n+3}$) generation.

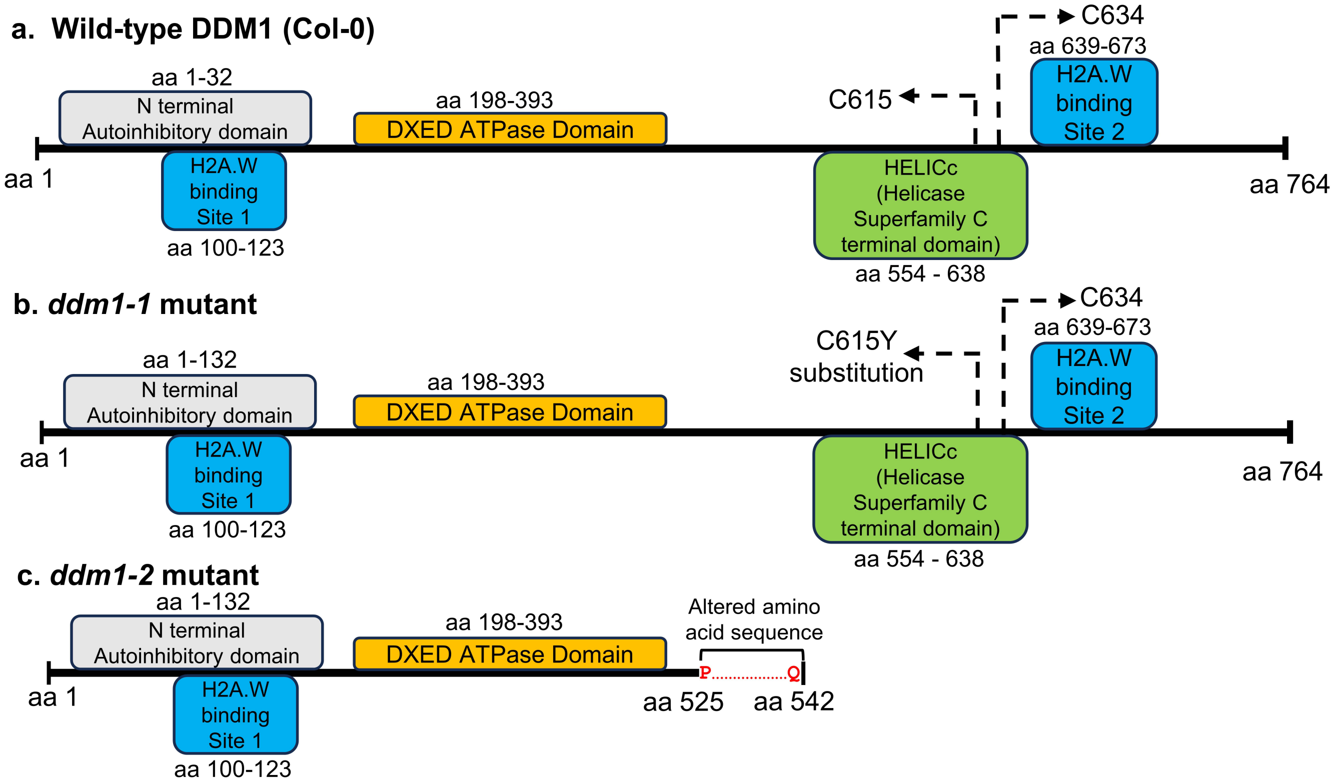


**Figure S12**: Diagrammatic representation of different domains of DDM1 protein. Cartoons show known functional protein domains in **(a)** WT DDM1 (Col-0), **(b)** *ddm1-1*, and **(c)** *ddm1-2* mutants.
aa=amino acid, H2A.W= Histone 2A variant W, C=cysteine, T=tyrosine
Functional domains are not drawn to scale with respect to number of amino acids.

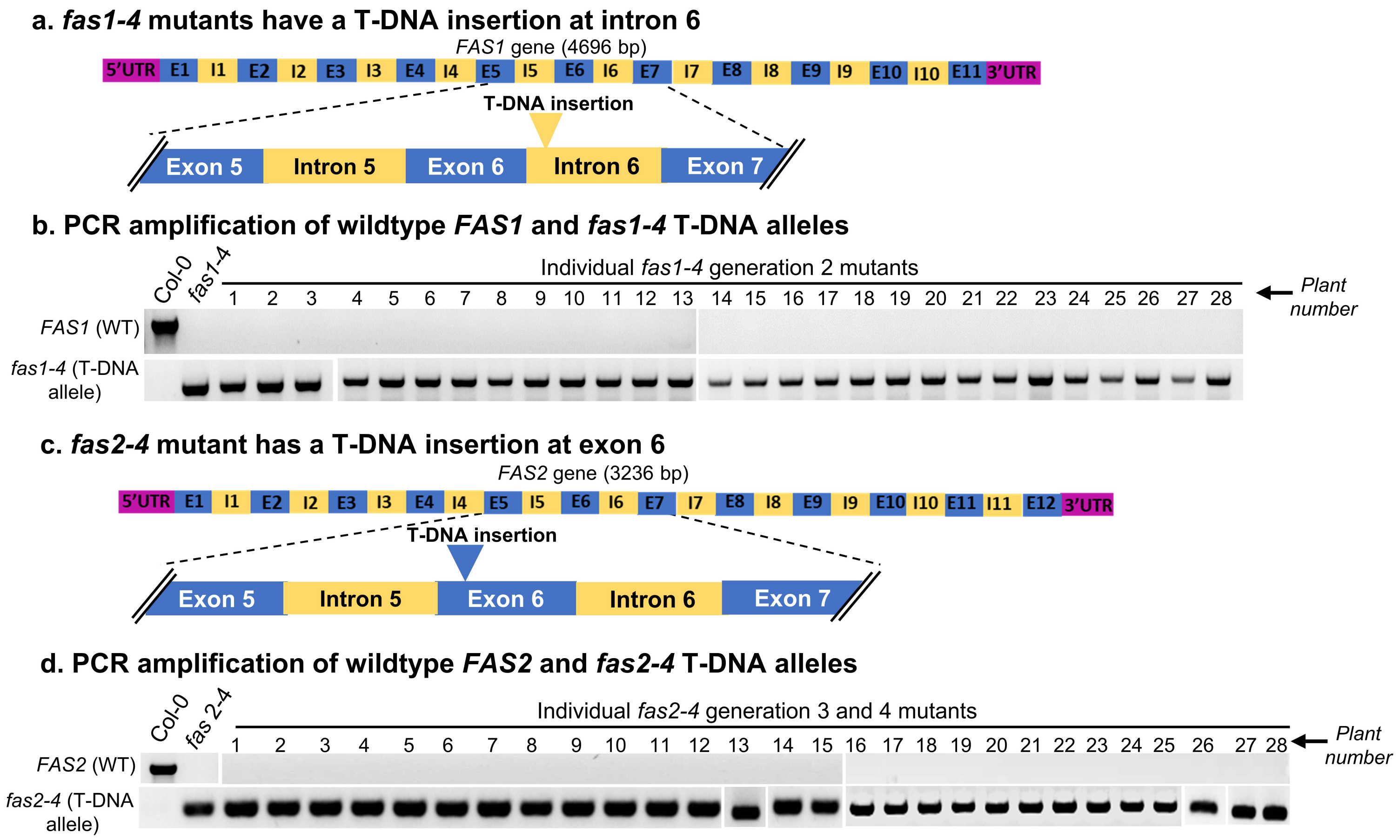


**Figure S13**: Locations of T-DNA insertion sites in *fas* mutants and their genotyping. **a** Cartoon shows the location of the T-DNA insertion site in *FAS1* gene in the *fas1-4* mutants. **b** Gel images show PCR amplification of the *FAS1* wildtype and T-DNA mutant allele in Col-0 and *fas1-4* mutants, respectively. **c** Cartoon shows the location of the T-DNA insertion site in the *FAS2* gene in the *fas2-4* mutants. **d** Gel images show PCR amplification of the *FAS2* wildtype and T-DNA mutant allele in Col-0 and *fas2-4* mutants, respectively.

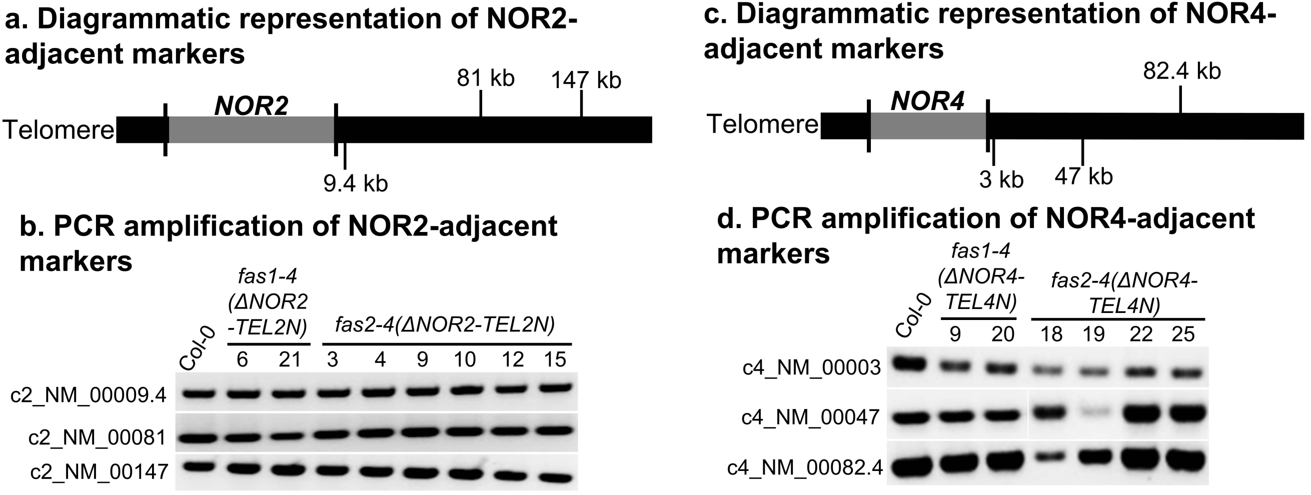


**Figure S14**: NOR-adjoining regions are intact in *fas1-4* and *fas2-4* mutants. **a** Cartoon depicts NOR2-adjacent markers. **b** The gel images show PCR amplification of NOR2-adjacent markers in WT Col-0, *fas1-4*, and *fas2-4* mutants. **c** Cartoon depicts NOR4-adjacent markers. **d** The gel images show PCR amplification of NOR4-adjacent markers in WT Col-0, *fas1-4*, and *fas2-4* mutants.

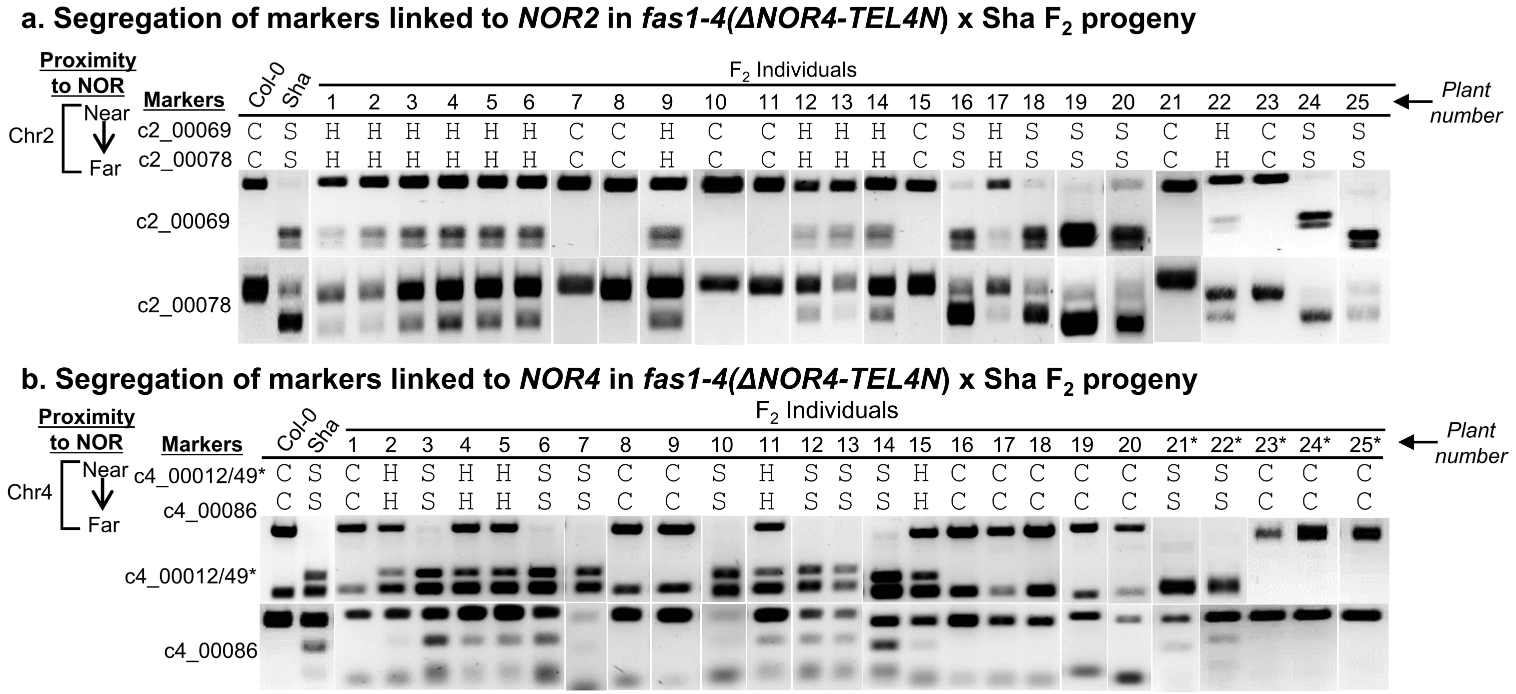


**Figure S15**: Genotyping of $F_2$ progenies of *fas1-4(ΔNOR4-TEL4N)* x Sha. The panels show gel images of length polymorphism-based PCR markers and CAPS markers that distinguish Col-0 and Sha alleles in the regions adjacent to *NOR2* and *NOR4* **a** Segregation of markers linked to *NOR2* in *fas1-4(ΔNOR4-TEL4N)* x Sha $F_2$ progenies. **b** Segregation of markers linked to *NOR4* in *fas1-4(ΔNOR4-TEL4N)* x Sha $F_2$ progenies.
C=Col-0, S=Sha, H=heterozygous.

### PCR amplification of *NOR2* markers N2-m1 and N2-m2

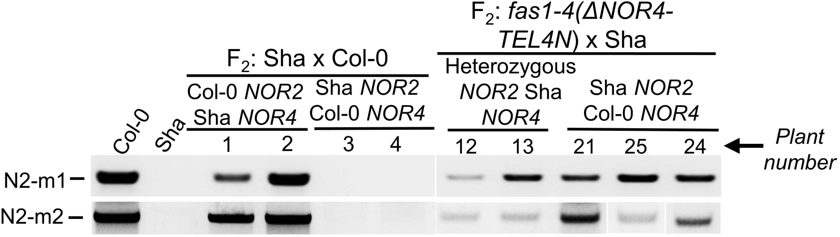


**Figure S16:** In *fas1-4*(*ΔNOR4-TEL4N*) mutants, telomere-proximal regions of *NOR4* rDNA and the associated telomeres are lost and replaced by the corresponding sequences from the *NOR2*. Gel images show PCR amplification of *NOR2* markers N2-m1 and N2-m2 in informative F$_2$ individuals of Sha x Col-0 and *fas1-4*(*ΔNOR4-TEL4N*) x Sha, and Col-0 and Sha controls.

**a. Chi-square test results for NOR2-linked marker for c2_00069 in $F_2$ progeny of *fas1-4(ΔNOR4-TEL4N)* x Sha**

| Genotype | Observed | Expected | χ2 | df | P (alpha) |
|---|---|---|---|---|---|
| Col-0 | 32 | 33 | 2.484 | 2 | Not significant |
| Heterozygous | 74 | 66 | | | |
| Sha | 26 | 33 | | | |
| Total number of plants genotyped: 132 | | | | | |

**b. Chi-square test results for NOR4-linked marker c4_NM_00047 in $F_2$ progeny of *fas1-4(ΔNOR4-TEL4N)* x Sha**

| Genotype | Observed | Expected | χ2 | df | P (alpha) |
|---|---|---|---|---|---|
| Col-0 | 21 | 21.25 | 0.0118 | 2 | Not significant |
| Heterozygous | 43 | 42.5 | | | |
| Sha | 21 | 21.25 | | | |
| Total number of plants genotyped: 85 | | | | | |

**Figure S17**: Chi-square statistic analyses show that NOR2- and NOR4-linked markers are segregating normally in the Mendelian fashion in $F_2$ progenies of *fas1-4*(*ΔNOR4-TEL4N*) x Sha. Tables show χ2 test results for **a** NOR2-linked marker c2_00069 and **b** NOR4-linked marker c4_00047 in the $F_2$ progeny of *fas1-4*(*ΔNOR4-TEL4N*) x Sha.
df=degrees of freedom; p= probability.

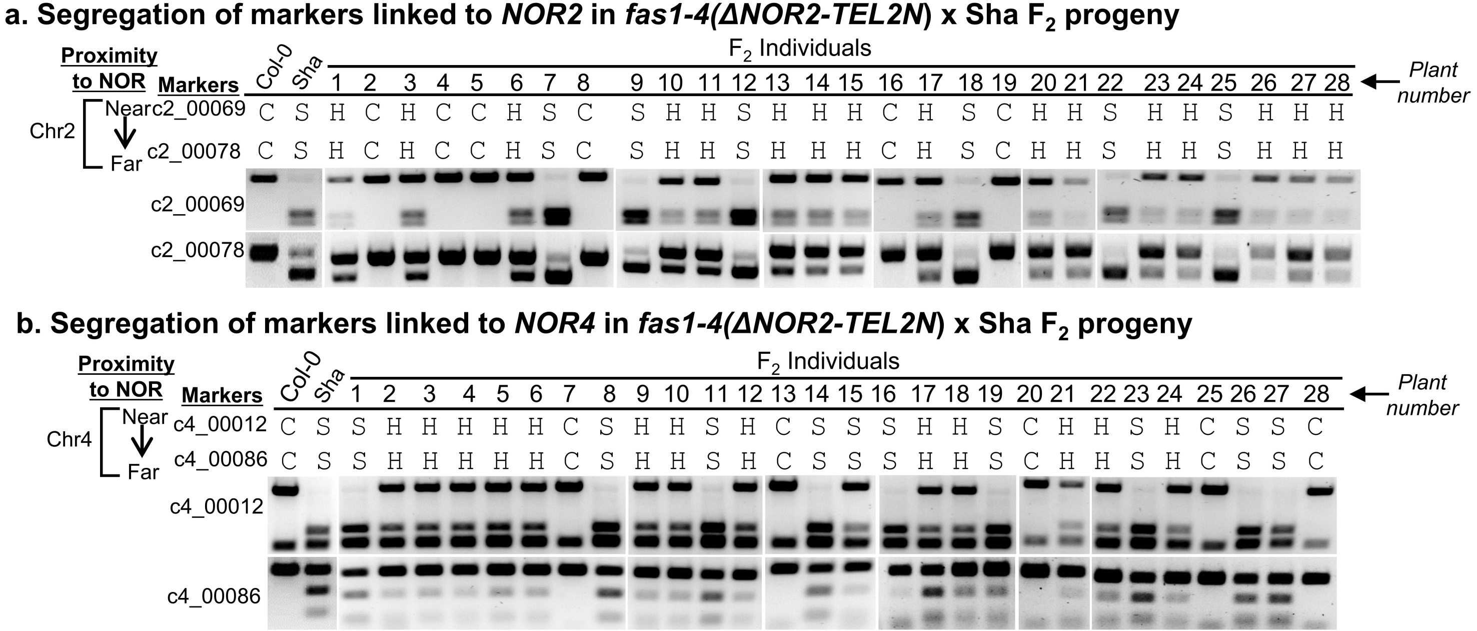


**Figure S18**: Genotyping of $F_2$ progenies of *fas1-4(ΔNOR2-TEL2N)* x Sha. The panels show gel images of length polymorphism-based PCR markers and CAPS markers that distinguish Col-0 and Sha alleles in the regions adjacent to *NOR2* and *NOR4.* **a** Segregation of markers linked to *NOR2* in *fas1-4(ΔNOR2-TEL2N)* x Sha $F_2$ progenies. **b** Segregation of markers linked to *NOR4* in *fas1-4(ΔNOR2-TEL2N)* x Sha $F_2$ progenies.
C=Col-0, S=Sha, H=heterozygous.

### PCR amplification of *NOR4* markers N4-m1 and N4-m2

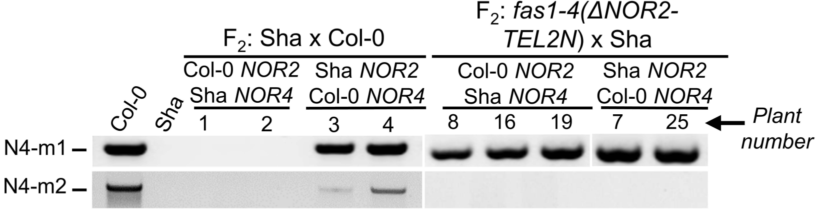


**Figure S19:** In *fas1-4*(*ΔNOR42-TEL2N*) mutants, telomere-proximal regions of *NOR2* rDNA and the associated telomeres are lost and replaced by the corresponding sequences from the *NOR4*. Gel images show PCR amplification of *NOR4* markers N4-m1 and N4-m2 in informative $F_2$ individuals of Sha x Col-0 and *fas1-4*(*ΔNOR2-TEL2N*) x Sha and Col-0 and Sha controls.

**a. Chi-square test results for NOR2-linked marker c2_00069 in $F_2$ progeny of *fas1-4 (ΔNOR2-TEL2N)* x Sha**

| Genotype | Observed | Expected | χ2 | df | P (alpha) |
|---|---|---|---|---|---|
| Col-0 | 17 | 28 | 5.910 | 2 | Not significant |
| Heterozygous | 65 | 56 | | | |
| Sha | 30 | 28 | | | |
| Total number of plants genotyped: 112 | | | | | |

**b. Chi-square test results for NOR4-linked marker c4_00086 in $F_2$ progeny of *fas1-4 (ΔNOR2-TEL2N)* x Sha**

| Genotype | Observed | Expected | χ2 | df | P (alpha) |
|---|---|---|---|---|---|
| Col-0 | 24 | 24.75 | 0.272 | 2 | Not significant |
| Heterozygous | 48 | 49.5 | | | |
| Sha | 27 | 24.75 | | | |
| Total number of plants genotyped: 99 | | | | | |

**Figure S20**: Chi-square statistic analyses show that NOR2- and NOR4-linked markers are segregating normally in the Mendelian fashion in $F_2$ progenies of *fas1-4*(*ΔNOR2-TEL2N*) x Sha. Tables show χ2 test results for **a** NOR2-linked marker c2_00069 and **b** NOR4-linked marker c4_NM_00086 in $F_2$ progeny of *fas1-4*(*ΔNOR2-TEL2N*) x Sha.
df=degrees of freedom; p= probability.

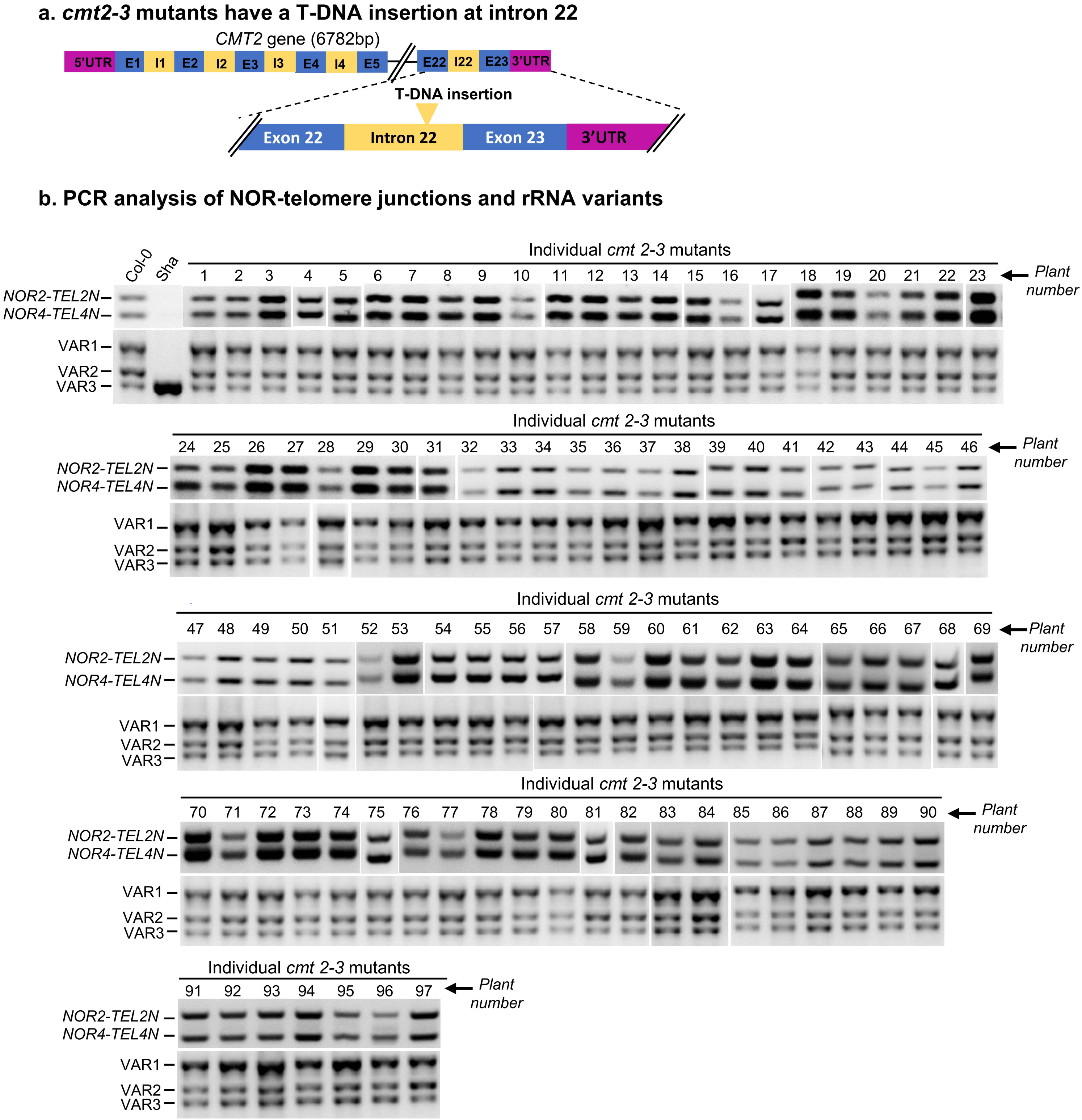


**Figure S21:** The *cmt2-3* mutants did not show any observable genomic instabilities of rDNA or the associated telomeres. **a** Cartoon shows the location of the T-DNA insertion site in the *CMT2* gene. **b** The panel of gel images show PCR analysis of NOR-telomere junctions and rRNA variants of *cmt2-3* mutants.

#### PCR amplification of wildtype *CMT2* and *cmt2-3* T-DNA alleles

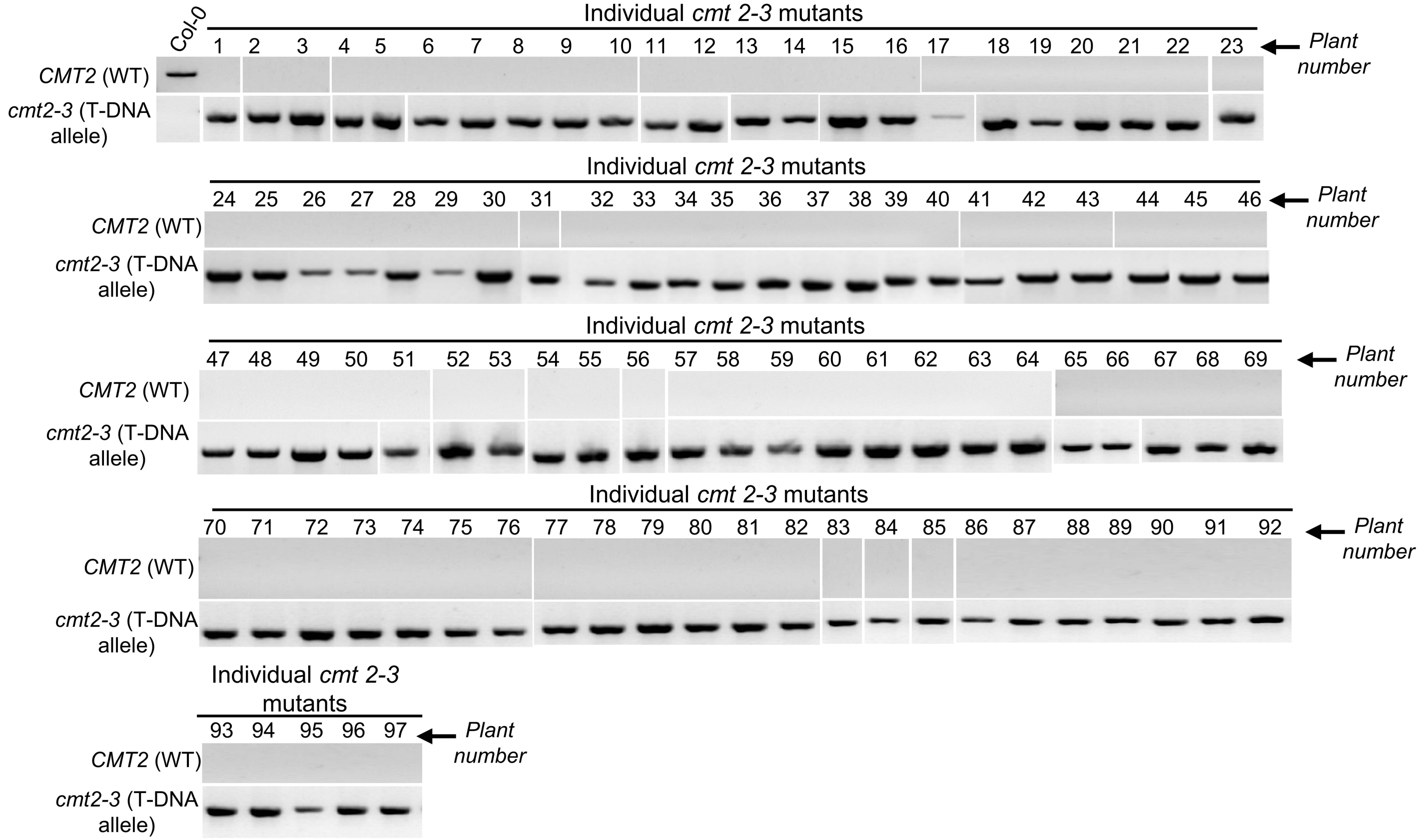


**Figure S22**: Genotyping of *cmt2-3* mutants. Gel images show PCR amplification of the *CMT2* wild-type and T-DNA mutant allele in Col-0 and *cmt2-3* mutants, respectively.

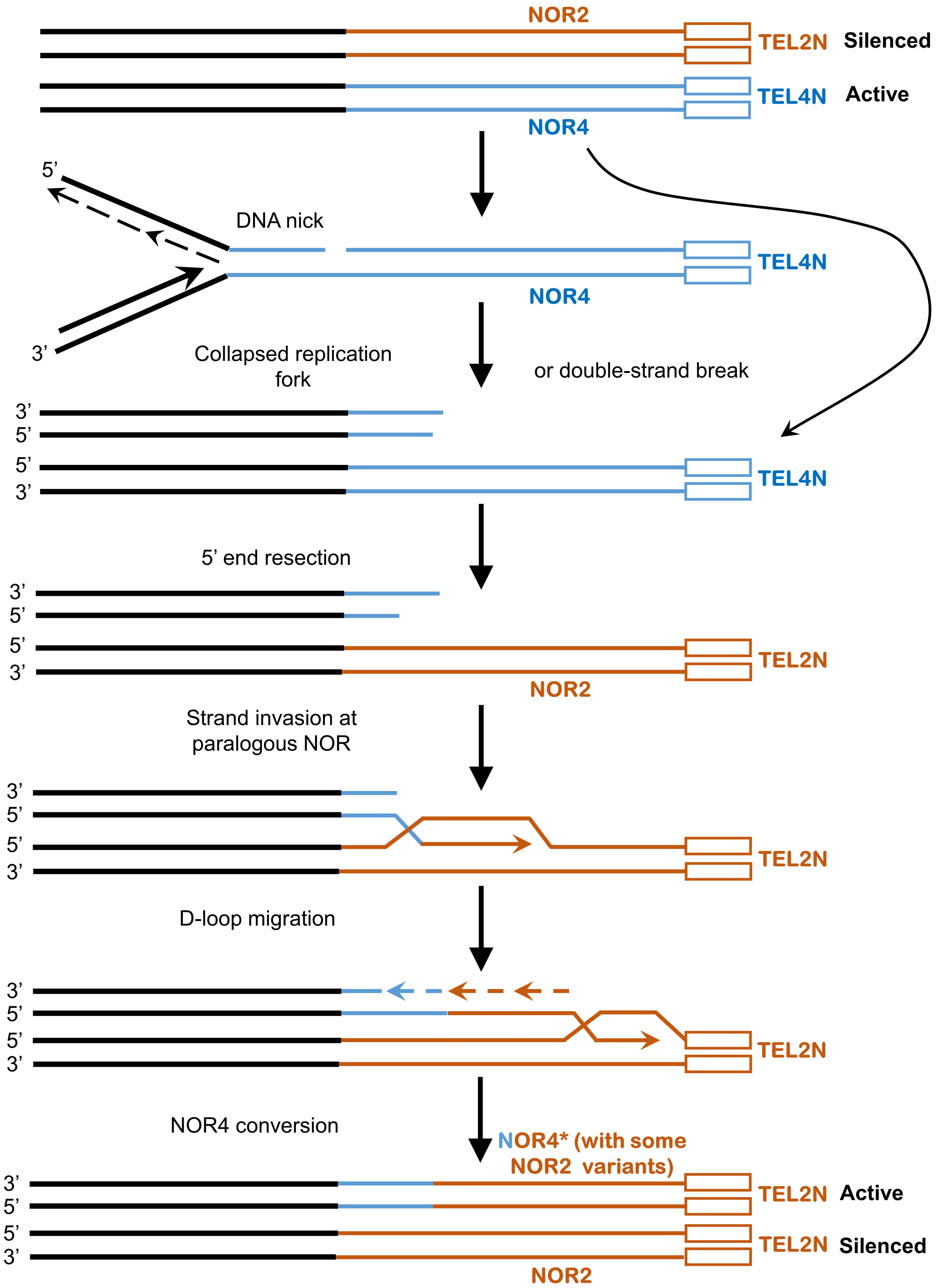


**Figure S23. Model for replacement of *NOR4-TEL4N* sequences by *NOR2-TEL2N* sequences via break-induced replication.** Within *NOR4*, a double-strand break, potentially initiated by a nick in the DNA template for the replicative lagging strand, initiates strand invasion of the nearly identical chromosomal sequences of *NOR2*. Repair by break-induced replication, extending to the end of the chromosome, converts *NOR4* and *TEL4N* to the corresponding sequences of *NOR2* and *TEL2N*. rRNA gene subtypes silenced at *NOR2* now escape silencing at converted *NOR4**. The diagram depicts proposed break-induced replication (BIR)-mediated *NOR4* conversion. We hypothesize same model for *NOR2* conversion also.
